# Post-Translational Modifications Soften Vimentin Intermediate Filaments

**DOI:** 10.1101/2020.06.05.135780

**Authors:** Julia Kraxner, Charlotta Lorenz, Julia Menzel, Iwan Parfentev, Ivan Silbern, Manuela Denz, Henning Urlaub, Blanche Schwappach, Sarah Köster

**Affiliations:** Institute for X-Ray Physics, University of Göttingen, 37077 Göttingen, Germany; Department of Molecular Biology, University Medical Center Göttingen, 37073 Göttingen, Germany; Bioanalytical Mass Spectrometry, Max Planck Institute for Biophysical Chemistry, 37077 Göttingen, Germany; Bioanalytic Group, Institute of Clinical Chemistry, University Medical Center Göttingen, 37075 Göttingen, Germany; Cluster of Excellence “Multiscale Bioimaging: from Molecular Machines to Networks of Excitable Cells” (MBExC), University of Göttingen, Germany; Max Planck School “Matter to Life”

## Abstract

The mechanical properties of biological cells are determined by the cytoskeleton, a composite biopolymer network consisting of microtubules, actin filaments and intermediate filaments (IFs). By differential expression of cytoskeletal proteins, modulation of the network architecture and interactions between the filaments, cell mechanics may be adapted to varying requirements on the cell. Here, we focus on the intermediate filament protein vimentin and introduce post-translational modifications as an additional, much faster mechanism for mechanical modulation. We study the impact of phosphorylation on filament mechanics by recording force-strain curves using optical traps. Partial phosphorylation softens the filaments. We show that binding of the protein 14–3–3 to phosphorylated vimentin IFs further enhances this effect and speculate that in the cell 14–3–3 may serve to preserve the softening and thereby the altered cell mechanics. We explain our observation by the additional charges introduced during phosphorylation.

---

The cytoskeleton of eukaryotes consists of three filament systems – microtubules, actin filaments and intermediate filaments (IFs) – which form a complex biopolymer network. The exact composition of the cytoskeleton and the interplay between the three filament types largely influence the mechanical properties of different cell types.^1^ Microtubules and actin filaments are highly conserved throughout cell types, whereas more than 70 human genes encode for IFs,^2^ yet they all share the same hierarchical assembly pathway from monomers to filaments. The secondary structure of the monomers consists of an *α*-helical rod domain, flanked by intrinsically disordered head and tail domains.^3,4^ During assembly, two monomers align and form parallel coiled-coil dimers, two dimers form antiparallel, half-staggered tetramers, and multiple tetramers constitute a unit-length filament (ULF) (see Fig. 1a). Subsequent longitudinal annealing yields elongated filaments with a diameter of about 10 nm.^5^ This hierarchical structure of IFs, in contrast to polar microtubules or actin filaments, grants IFs their unique mechanical properties.^6–8^

**Figure 1:**
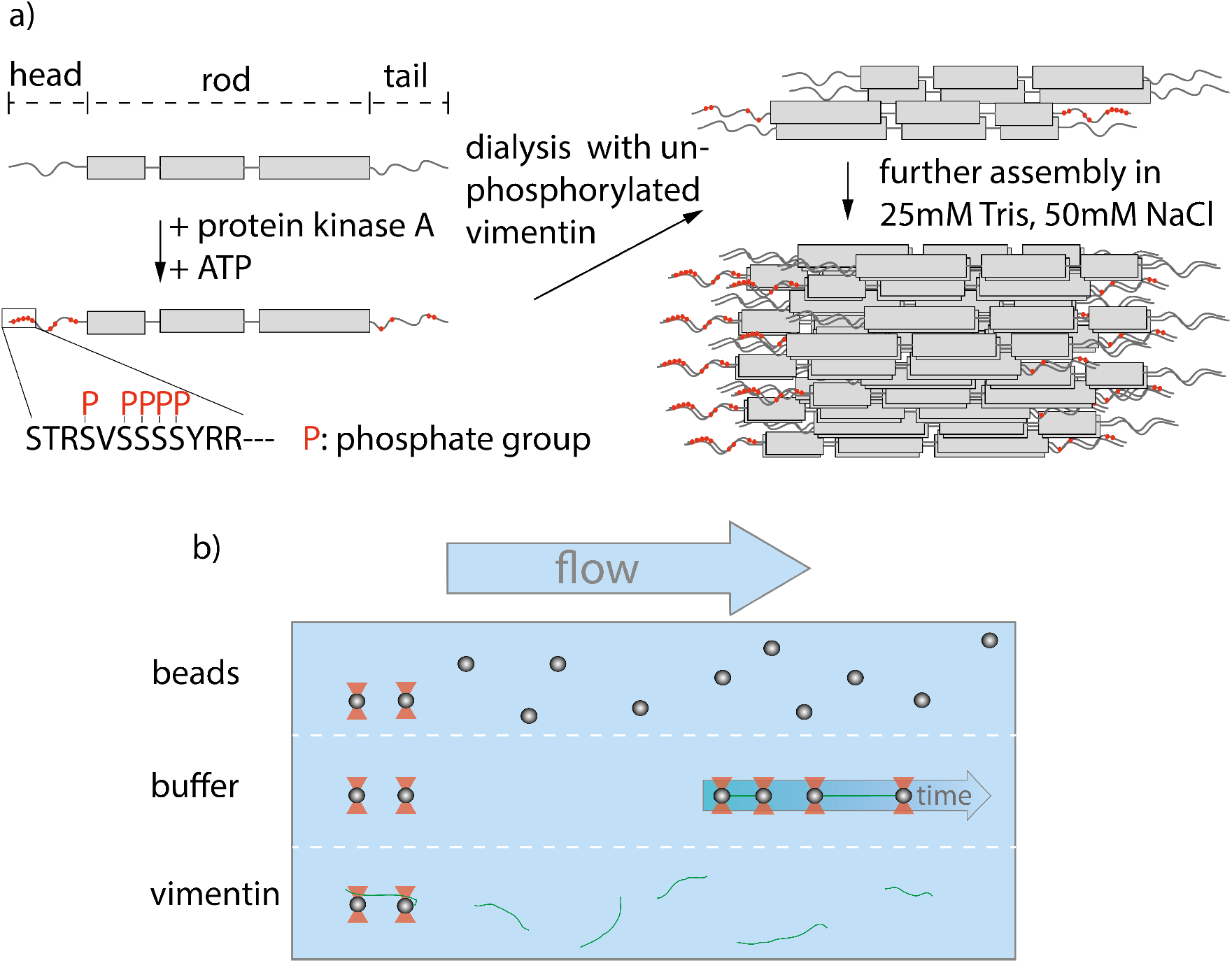
Experimental design and setup. a) Assembly process of partially phosphorylated vimentin. Vimentin monomers consist of a central rod domain flanked by intrinsically disordered head and tail domains. A part of the monomers are phosphorylated by addition of protein-kinase A and ATP leading to several phosphorylated sites in the head and tail, highlighted in red. The phosphorylated monomers are mixed with unphosphorylated vimentin prior to assembly by dialysis into a salt-containing buffer. b) Overview of the microfluidic chip used within the optical trap. The different compartments are realized by laminar flow and the measurements are performed in the buffer region.

The most abundant IF protein, vimentin, is found in cells of mesenchymal origin.^5,9^ Single vimentin filaments are highly extensible and can be elongated up to at least 4.5 fold.^7,10,11^ During elongation, three regimes are observed in the force-strain curves: an initial linear (elastic) increase, a plateau region and a subsequent stiffening.^6,7,12,13^ These different stretching regimes have been linked to structural changes in the rod domain of IFs^12^ such as the opening of *α* helices during the plateau regime.^6,14,15^ Additionally, vimentin filaments are able to dissipate large amounts of the input energy while being stretched. ^14^ Consequently, vimentin IFs act as shock-absorbers for mechanical protection of the cell as well as scaffolds that help to maintain cell shape and organization of the cytoskeleton. ^9,16^

An effective way to tune the mechanics of IFs is the variation of the charges of the amino acids constituting the protein.^17^ One cellular mechanism for such charge variation are post-translational modifications (PTMs). Within the IF cytoskeleton, several types of PTMs have been described^18,19^ and the most abundantly studied PTM in IFs is phosphorylation, which is involved in regulation of IF dynamics by leading to disassembly and in providing binding sites to signaling proteins.^20,21^ It has been shown that the phosphorylation of vimentin by protein-kinase A (PKA) leads to various phosphorylated sites, most of which are positioned in the head region (see Fig. 1a).^21^ The importance of the head domain in the assembly process was shown in Refs. 3,22, stressing the obvious link between phosphorylation and assembly dynamics. Although such changes in the molecular interactions during assembly are likely to influence the behavior of the fully assembled filaments, the influence of phosphorylations on filament mechanics is not yet resolved.

A further interesting aspect of phosphorylation is the ability of certain proteins to bind to the modified sites. One such protein is 14–3–3,^23^ which is involved in several cellular processes like signal transduction, adhesion and inhibition of tumorigenesis.^24^ The role of this protein depends on the interaction partner, *e.g*. it binds to keratin during the cell cycle^25^ and affects the assembly dynamics of neurofilaments.^26^ However, the role of 14–3–3 for vimentin is unknown.

Here we investigate the effect of phosphorylation and 14–3–3 on vimentin mechanics by studying precise force-strain curves from optical trap experiments. We find that the filaments soften with increasing amount of phosphorylated protein within the filament and that interaction with the protein 14–3–3 further enhances this effect. By combining our mechanical measurements with mass spectrometry, cross-linking, phosphomimicry and numeric modeling, we are able to attribute the softening to reduced lateral coupling of monomers within the filaments.

To investigate whether phosphorylation of vimentin influences filament mechanics, we record force-strain curves using optical traps. As complete phosphorylation of vimentin filaments leads to disassembly, ^21^ we perform the stretching experiments on partially phosphorylated vimentin filaments with varying percentages of 1, 5 or 10%, as sketched in Fig. 1a. The incorporation of the phosphorylated vimentin is confirmed by sodium dodecyl sulfatepolyacrylamide gel electrophoresis (SDS-page) and by fluorescence microscopy, shown in Supporting Figs. S1 and S2. The single filaments are covalently bound to two polystyrene beads and each bead is captured in an optical trap for precise force measurements. Filaments with 10% phosphorylated protein incorporated assemble to the same total length as unphophorylated filaments, see Supporting Fig. S3. Additionally, we determine the distance between both beads and calculate the strain as 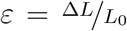, where *L*_0_ is the original filament length and Δ*L* the additional extension upon application of a force *F*. We perform all measurements in the “buffer” channel, *i.e*. without any disturbance from surrounding beads or protein, see Fig. 1b. During stretching, we observe the characteristic response of vimentin to the force, including the linear increase for small strains, the plateau region, where the *α* helices open up^6,7,14^ and the subsequent stiffening of the filaments. We perform all measurements in 25 mM Tris-HCl, 50 mM NaCl, pH 7.5 and the curves are comparable to the ones previously recorded in phosphate buffer, ^7,14^ see Supporting Fig. S4. The pH of the measuring buffer is chosen such that it is comparable to most *in vitro* studies on vimentin filaments. For a lower pH, we expect the same shape of the curve, but slightly shifted to higher forces, as shown recently in Ref. 17. Such a typical curve for our standard condition is shown in Fig. 2a, green. We show an average curve here for clarity, however, the individual data sets are shown in the Supporting Fig. S5. The measurements end when the force becomes too large for the optical traps, *i.e*. at roughly 550-600 pN. Even though strong phosphorylation leads to disassembly of vimentin filaments, ^21^ we do not observe an increase in the number of breakage events upon increasing phosphorylation. Fig. 2a also shows corresponding data for filaments containing a certain percentage of phosphorylated protein (shades of blue, for color code see legend). To quantify the force-strain data, we focus on the Young’s modulus, which we calculate from the initial slope in the linear regime up to a force of 130 pN of each curve, as a measure of the filament stiffness. The Young’s modulus for the unphosphorylated vimentin filaments is in good agreement with the results found in Ref. 7. The analysis procedure is visualized in Supporting Fig. S6. The Young’s moduli plotted in Fig. 2d (green) show a strong decrease with increasing percentage of phosphorylated protein. Further analysis of the force-strain curves shows a decrease of the force at the onset of the plateau and an increase of the maximum strain reached with an increasing amount of phosphorylation, shown in Supporting Fig. S7.

**Figure 2:**
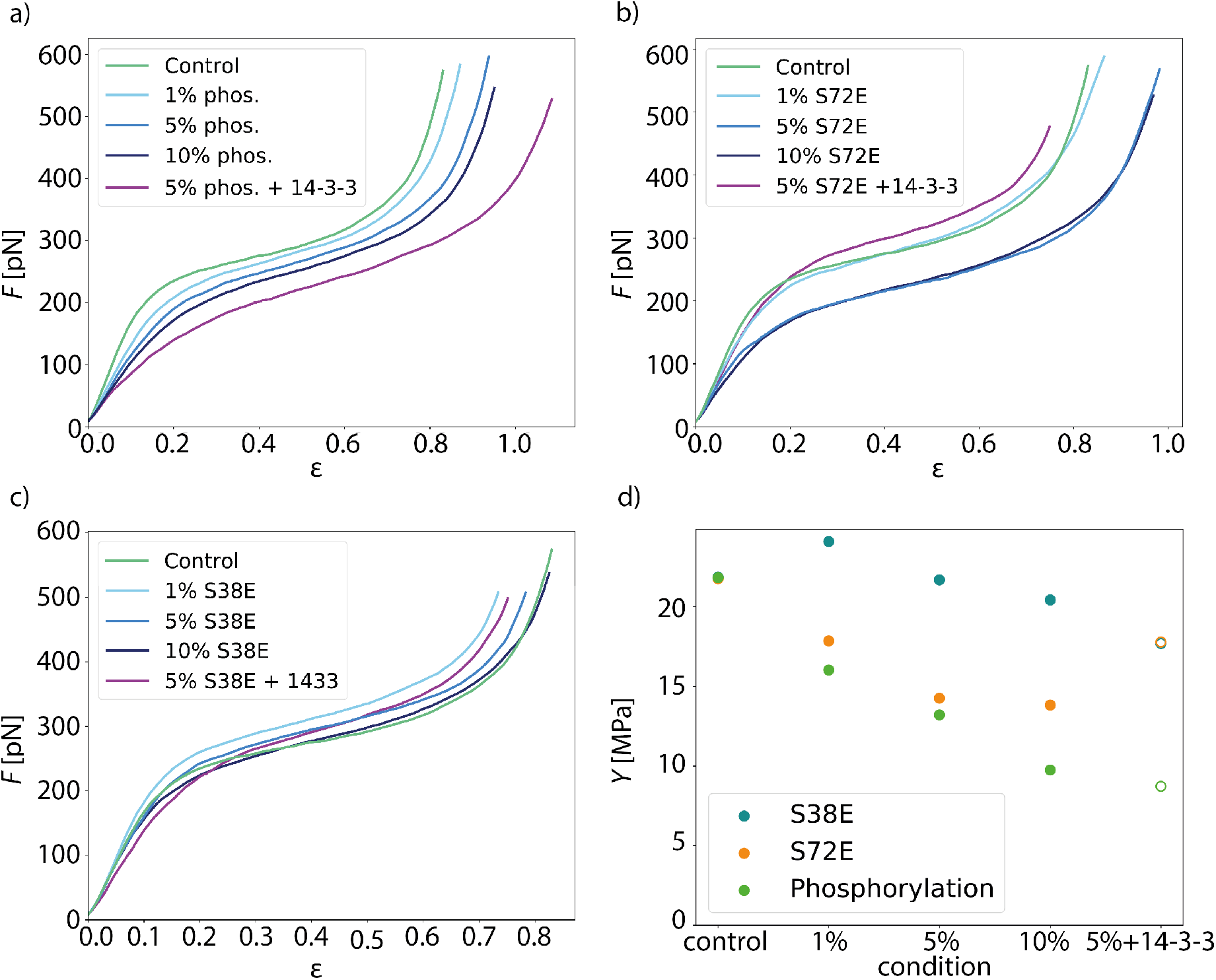
Additional negatively charged amino acids soften vimentin filaments. a) Mean force-strain curves for partially phosphorylated filaments: control (unphosphorylated, green), 1% phosphorylation (light blue), 5% phosphorylation (medium blue), 10% phosphorylation (dark blue) and 5% phosphorylation with 14–3–3 (magenta). With increasing amount of phosphorylated vimentin incorporated, the filaments become softer; the effect is even more pronounced in the presence of the protein 14–3–3. b, c) Mean force-strain curves for the phosphomimicry data. The color code for the individual conditions is the same as in a. b) The mean curves for the phosphomimetic mutant S72E show a similar trend as the phosphorylation data except for the filaments incubated with 14–3–3. c) The mean curves of the phosphomimetic mutant S38E do not show a systematic softening regardless of whether the filaments were incubated with 14–3–3 or not. d) Comparison of the different data sets. The Young’s modulus *Y*, which is a measure of the filament stiffness, is shown in dependence of the amount of phosphorylated or phosphomimetic protein. The phosphorylation (green) and the S72E data (orange) show a softening with increasing phosphorylation or phosphomimicry, whereas the S38E data (blue) remain fairly constant. The Young’s moduli for filaments with additional 14–3–3 are depicted as open symbols, as these are only an estimate using the radius of the 5% condition instead of the actual radius that cannot be measured by SAXS.

Upon phosphorylation of vimentin by PKA, several amino acids are modified. To determine these sites, we perform an LC-MS (liquid chromatography mass spectrometry) analysis. Fig 3a shows that phosphorylated sites are dispersed throughout the whole protein, but the most abundant ones (log_2_(I) > 0, red lines) are all found in the head region of vimentin. This is in agreement with the head domain of vimentin being a serine rich region and containing multiple motives required for kinases. ^27^ Strikingly, the positions S71 and S72 are always phosphorylated simultaneously and show the highest degree of phosphorylation. Hence, we speculate that these two positions are the main phosphorylation sites, whereas all other positions occur only sporadically and therefore differ from monomer to monomer. We employ phosphomimicry to investigate the effect of defined phosphorylated sites and choose two of the most abundantly phosphorylated sites that also occur *in vivo*, S38 and S72. ^21^ We perform the same force-strain measurements as described above. For the mutant S72E, we observe a similar trend as in the phosphorylation data including enhanced softening, see Fig. 2b, green to dark blue. On the contrary, Fig. 2c, green to dark blue, shows filaments containing the mutation S38E and no systematic trend is observed. The individual data sets are provided in Supporting Figs. S8 and S9. When comparing these three different conditions, *i.e*. phosphorylation, mutation S72E and mutation S38E, we observe that the Young’s modulus, and therefore the filament stiffness, decreases with an increasing amount of phosphorylation (green) and mutation S72E (orange) but stays fairly constant for the mutation S38E (blue), see Fig. 2d. These results suggest that the phosphorylation at position S72 influences the filament mechanics whereas at position S38 it does not have any effect. As the filament radius is a critical parameter for calculation of the Young’s modulus, we perform small angle X-ray scattering (SAXS) experiments to determine the radius of gyration of the cross-section *R_c_*, see Fig. 3c-e. No changes in radius are observed. The individual SAXS curves are shown in Supporting Figs. S10-S12.

**Figure 3:**
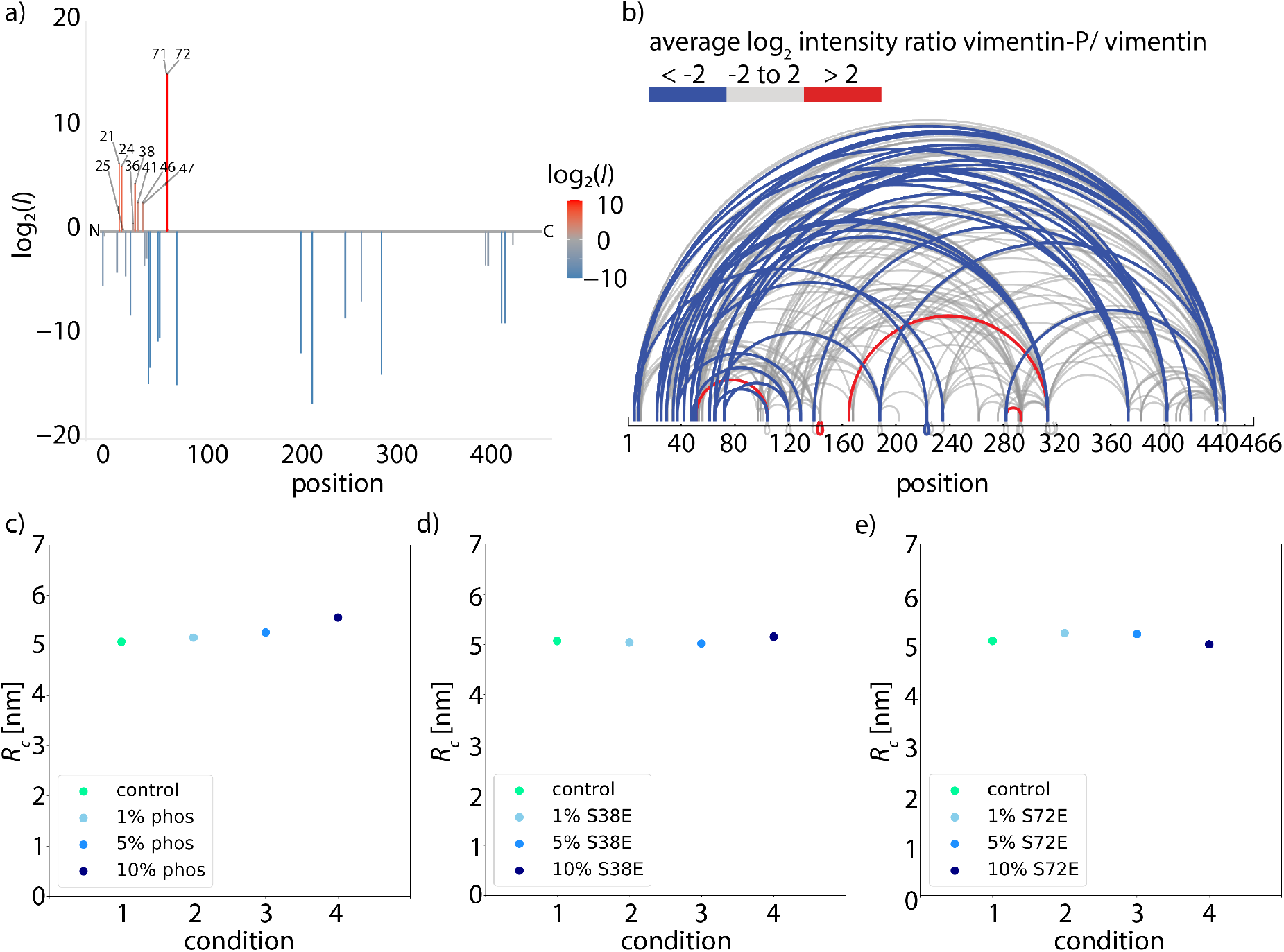
Structural changes caused by phosphorylation. a) Determination of phosphorylated sites in vimentin. Phosphorylated peptides are determined by LC-MS and the degree of phosphorylation of all identified sites is plotted using the log_2_ ratios of the LC-MS intensities of phosphorylated over unphosphorylated peptides (intensities *I*). Identified phosphorylation sites and the numbers of amino acids are listed. The color code represents the degree of phosphorylation of all identified phosphorylation sites. b) Mass spectrometry analysis of cross-linked tetramers. The ratio of the amount of cross-links found in the phosphorylated condition and the unphosphorylated condition is calculated. Linked amino acid positions within a tetramer are indicated by loops and the ratio of cross-links between the phosphorylated and unphosphorylated vimentin tetramers is shown by the color code. Blue loops show a decreased amount of cross-links in the phosphorylated condition, gray loops show no change in the ratio of cross-links between phosphorylated and unphosphorylated condition and red loops show an increase of cross-links in the phosphorylated condition. c)-e) SAXS experiments to analyze the influence of phosphorylation or phosphomimicry on the filament radius. c) Radius of gyration of the cross-section *R_c_* for the different conditions with the control measurement (green), 1% phosphorylation (light blue), 5% phosphorylation (medium blue) and 10% phosphorylation (dark blue). The radius shows no change for the different conditions. d) Radius of gyration *R_c_* for the different conditions with the control measurement (green), 1% S38E (light blue), 5% S38E (medium blue) and 10% S38E (dark blue). The radius shows no change for the different conditions. e) Radius of gyration *R_c_* for the different conditions with the control measurement (green), 1% S72E (light blue), 5% S72E (medium blue) and 10% S72E (dark blue). The radius shows no change for the different conditions.

To explain the softening of vimentin filaments with increasing amount of phosphorylation, we consider previous studies that have shown neighboring dimers to be coupled by electrostatic interactions between specific positively charged amino acids in the head, namely R23, R28, R36 and R45, and the negatively charged coiled-coils^3,28,29^ as sketched in Fig. 4b. When vimentin becomes phosphorylated, the positive charges of the head domain are flanked by negative charges of the phosphorylated amino acids as shown in Fig. 4a, which diminishes the electrostatic attraction between the head and the coiled-coils. This observation raises the question of whether the shift in filament stiffness can be explained by weaker coupling. Therefore, we run Monte-Carlo simulations to understand how the coupling of dimers affects the force-strain curves of vimentin by extending the model from Refs. 8,14 as described in detail in the SI. We model a vimentin monomer by including the *α* helices in the rod domain as the spring constant *κ_α_* and as elements, which extend in length at a certain force, thereby transitioning to an unfolded state *u*.^8,14,15^ The model in Ref. 8 assumes a strong coupling *within* dimers, tetramers or any other subunit of the ULF and weaker coupling *between* these subunits. In that work, the force-strain curves of filaments with smaller coupled subunits exhibit a lower plateau than the force-strain curves of filaments with larger coupled subunits. As demonstrated in Ref. 17, the model can also explain different maximum strains of filaments in different conditions either by a different number of *α* helices which unfold or a different extension of single *α* helices. Yet, this previous model does not explain a pronounced decrease in Young’s modulus as observed here.

**Figure 4:**
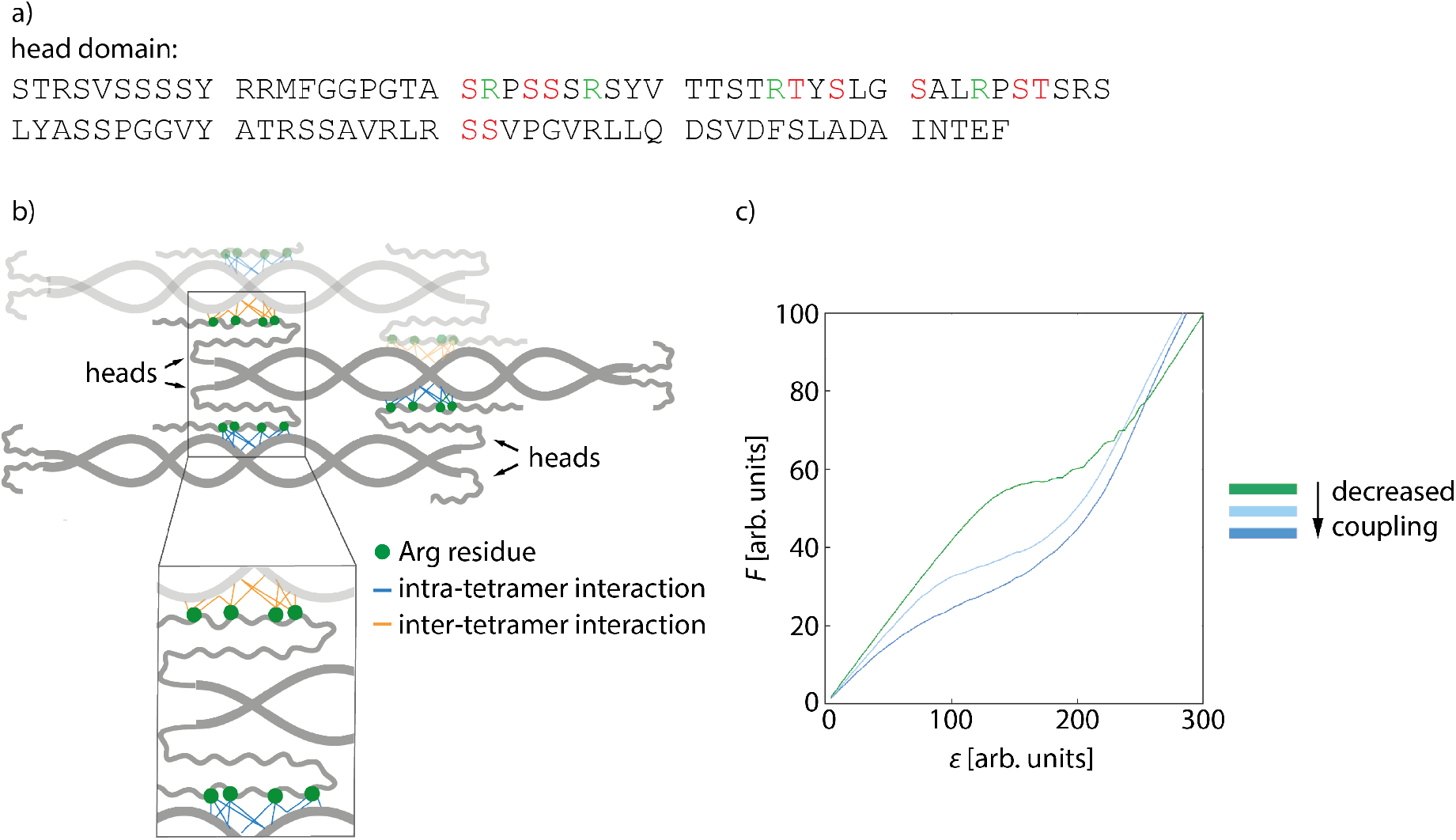
Softening of phosphorylated filaments is due to decreased lateral interactions. a) The amino acid sequence of the head domain of vimentin with positively charged amino acids (green) and phosphorylated sites (red). b) Sketch of a vimentin tetramer. The positively charged amino acids in the head domain (green dots) interact with the negatively charged coils of the neighboring dimer. Intra-tetramer interactions are depicted in blue, the neighboring tetramer is illustrated in light gray and inter-tetramer interactions are shown in orange. c) Results of numerical model for the softening of the filaments due to decreased lateral coupling. The influence of decreased coupling is shown in the force-strain plot where the completely coupled system is shown in green and the lateral coupling is decreased from light blue to dark blue.

We thus supplement the model from Ref. 8 by spring constants *κ_bt_*, which represent bonds between tetramers, and *κ_bd_* for bonds between dimers within the tetramer. In case of more phosphorylated monomers in the filament, the filament subunits do not interact as strongly as without phosphorylated monomers because the additional negative charges repel each other. Thus, not all bonds between tetramers and dimers can form, so that the spring constants *κ_bd_* and *κ_bt_* do not contribute to the overall spring constant and the filaments get softer as shown for the simulated force-strain curves in Fig. 4d. Consequently, the initial slope of the force-strain curves decreases with more phosphorylated monomers as shown in Fig. 2c. Our results show that the negative charge at position S72, in particular, has an effect on filament mechanics: the phosphomimetic mutant shows a very similar trend compared to the phosphorylation data (Fig. 2d). Indeed, this position is also found to be the mayor phosphorylation site (see Fig. 3a). By contrast, an additional negative charge at position S38 has no effect on the filament mechanics. Therefore, we conclude that a decreased coupling around position S72 is crucial for the shift in mechanics whereas a decreased coupling at position S38 has no effect.

We confirm these numerical findings with mass spectrometry cross-linking experiments. When comparing cross-linked phosphorylated vimentin tetramers and unphosphorylated vimentin tetramers in Fig. 3b, fewer cross-links are found in the phosphorylated state, blue lines. All individual cross-linking positions can be found in Supporting Fig. S13. This supports our proposal that there are decreased interactions between neighboring dimers in phosphorylated vimentin which indicates that the lateral coupling of dimers is reduced. Despite these decreased interactions, we do not observe an increase in filament radius for increasing amount of phosphorylation, see Fig. 3c-e. We speculate that the softening effect due to phosphorylation is reversible when the phosphate group is removed again by phosphatases and filaments would recover their initial stiffness.

Besides the fact that phosphorylation modifies the protein itself, it also creates binding sites for other proteins, such as 14–3–3 in the case of phosphorylated vimentin.^23^ This raises the question of whether the binding of the protein 14–3–3 to phosphorylated vimentin also has an effect on the mechanics and we perform optical trap measurements of the vimentin/14– 3–3 complex. We confirm the interaction between vimentin and 14–3–3 by performing a streptavidin pulldown assay as shown in Supporting Fig. S14. Fig. 2d, open green circle, shows that for phosphorylated filaments the interaction with 14–3–3 softens the filaments even more. By contrast, the original stiffness of the unphosphorylated filaments is recovered for the mutant S72E, Fig. 2d, open orange circle, and we observe only a slight influence of 14–3–3 for the mutant S38E, Fig. 2d, open blue circle. The individual curves for these experiments are provided in Supporting Fig. S15. We show that 14–3–3 binds to the phosphorylated sites in the head domain of vimentin, as confirmed by cross-linking the complex and analyzing it with mass spectrometry, see Supporting Fig. S16. Assumedly, because of this interaction and the similar size of 14–3–3 compared to vimentin, it forces vimentin subunits further apart and thus enhances the decoupling and softening of vimentin filaments. Mostly, vimentin is cross-linked to position 78 of the amino acid sequence of 14–3–3 as shown in Supporting Fig. S16. By contrast, we cannot unambiguously determine the cross-linking position in the amino acid sequence of vimentin. However, our data show that the amino acids S38 and S72 in vimentin are not the binding sites for 14–3–3 as the complex of the two proteins is retrieved after phosphorylation of the phosphomimetic mutants as confirmed by cross-linking the complex, see Supporting Fig. S17. Additionally, we find that no complex is formed between 14–3–3 and the phosphomimetic mutants S38E and S72E. The corresponding SDS gel is shown in Supporting Fig. S17. Despite this lack of complex formation between 14–3–3 and the phosphomimetic mutant S72E, we observe a stiffening of the filaments upon addition of 14–3–3, indicating some kind of interaction between the proteins, such as electrostatic interactions. Such electrostatic interaction could possibly diminish the effect of the negative charges and thus lead to a recovery of the initial stiffness. Taking these results together, we propose that 14–3–3 protects the protein from phosphatases by binding to strategic phosphorylated sites as suggested in Ref. 30. Due to the large size of the protein 14–3–3, it sterically hinders the phosphatases, which can reverse the effect of the protein kinases, from binding to neighboring amino acid positions within the vimentin monomer. Thereby the protein 14–3–3 could be able to keep the vimentin filaments in the soft state for extended times, which might be important for *in vivo* situations.

Previous studies have shown that increased vimentin phosphorylation is required for efficient cellular migration^31^ and that it is relevant in metastasis^32,33^ which may be linked to a softer vimentin network and therefore render the cells more deformable. In general, phos-phorylation controls the assembly and disassembly dynamics of vimentin^21^ and in particular the phosphorylation of vimentin leads to disassembly. Phosphatase activity enables recovery to assembled filaments. If 14–3–3 binds to phosphorylated vimentin and inhibits the phosphatase activity, vimentin remains in the phosphorylated state. Such an effect would slow down the assembly dynamics, which is, however, crucial for cell adhesion, migration and signaling.^34^ In addition, the vimentin/14–3–3 complex builds a larger ensemble with the phosphorylated protein beclin1, which then promotes tumorigenesis.^35^ It was already suggested that vimentin might be a key regulator of tumorigenic pathways as this complex, namely 14–3–3, vimentin and beclin1, might prevent the dephosphorylation of the proteins within the ensemble and thereby inhibit antitumor activity in cells. ^36^

To conclude, we directly show how post-translational modifications, *i.e*. phosphorylation, change the mechanical properties of vimentin filaments: Vimentin filaments become softer with increasing amount of phosphorylated protein within the filament. These findings may help to understand the relation between the role of phosphorylation in cancer metastasis and the pronounced motility of metastasizing cells. The interaction of phosphorylated vimentin with 14–3–3 enhances this softening effect and may even protect the softer state. We suggest that these changes are induced by reduced electrostatic coupling within the ULF due to additional negative charges introduced by the phosphate groups and support this assumption by a physical model. We thus hypothesize that cells are able to fine-tune and adapt their mechanical properties locally and within seconds by modifications like phosphorylation according to specific external requirements.

## Material and Methods

An extended version of the Methods section can be found in the Supporting Information.

### Vimentin purification

Purification of recombinant human vimentin C328A with additional amino acids glycine-glycine-cystein at the C-terminus based on Herrmann *et al*.^29^ was performed.^14^ The additional amino acids were included to enable binding of the vimentin filaments to maleimide-functionalized beads and for fluorescent labeling of the monomers. Removing the cystein from the rod domain of the protein enables smooth assembly even after labeling. As we did not add zinc ions in our experiments,^37^ the missing cystein is not expected to influence the assembly and architecture of the filaments. Indeed, it has been shown that vimentin C328A assembles in the same way as wildtype vimentin.^38^ Two phosphomimetic mutants S38E and S72E of this vimentin C328A were produced according to the same protocol. We performed all control measurements with vimentin C328A and thus ensured that all observations were due to phosphorylation or phosphomimicry.

### Vimentin filament preparation and bead functionalization

Vimentin was fluorescently labeled with ATTO647N *via* maleimide chemistry^7^ to enable visualization of the filaments during the optical trapping experiments. To prepare the protein for filament assembly, the protein in storage buffer (8M urea, 5mM Tris-HCl, 1mM EDTA, 0.1mM EGTA, 0.01mM MAC and 250mM KCl, pH 7.5) was dialyzed against 6M urea, 5mM Tris-HCl at pH 8.4 and a stepwise dialysis against 5 mM Tris-HCl, pH 8.4 was performed. To initiate the assembly, the protein was dialyzed against 25mM Tris-HCl, pH 7.5 with 50mM NaCl over night. The final amount of labeled monomers within the filaments was about 4%. In analogy to the assembly of phosphorylated vimentin, the phosphomimetic mutants were mixed with vimentin C328A at mixing ratios of 1%, 5% and 10%. The mixing was performed in 8 M urea, *i.e*. prior to the dialysis step to ensure mixing at the monomer state of vimentin. Carboxylated polystyrene beads (Kisker Biotech GmbH & Co. KG, Steinfurt, Germany) with a diameter of 4.4 *μ*m were functionalized with maleimide according to Ref. 39.

### Phosphorylation of vimentin

We phosphorylated vimentin tetramers in 5mM Tris-HCl, pH 8.4 using cAMP-dependent protein kinase A (New England Biolabs, Frankfurt, Germany). The phosphorylated vimentin was mixed at the desired ratios of 1, 5 and 10% with unphosphorylated vimentin. Subsequently, a stepwise dialysis and the assembly were performed as described above.

### Optical trap measurements and analysis

For all measurements a commercial optical trap setup (C-trap, Lumicks, Netherlands), including a microfluidic chip and confocal microscopy, was used. For each measurement a fresh pair of beads was captured in the bead channel (see Fig.1b). The calibration of the optical trap was performed by analysis of the power spectral density of the thermal fluctuations of the trapped beads in the buffer channel. Afterwards they were moved to the vimentin filament channel, while scanning with the confocal microscope, until a single vimentin filament was captured on one bead. Then, the beads were moved back to the buffer channel where the filament was attached to the second bead. We ensured that the filament was relaxed to avoid prestrain. The measurements were performed by moving one bead at about 0.7 *μ*m/s to stretch the filament until it ruptured or the forces on the second bead were higher than the maximum trapping force. During the stretching, force-distance curves were recorded.

Data were analyzed with self-written Python codes. The calculation of the mean curves for each conditions was adapted from Ref. 8. For the calculation of the Young’s modulus, a linear fit up to a force of 130 pN was performed for the initial slope in the linear regime of the mean force-strain curves. The Young’s modulus was calculated *via* the ratio of stress and strain, 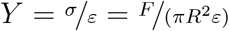. For the radius R, the corresponding values for the radius of gyration of the cross-section, R_*c*_, obtained from the small-angle X-ray scattering experiments, were used.

### Model

To simulate the force-strain behavior of a vimentin IF, we calculated all spring constants of the modeled elements and transition rates of possible reactions and ran a Monte-Carlo simulation with a self-written Matlab code (MathWorks, Natick, Massachusetts, USA) as in Ref. 8,14.

## Acknowledgement

The authors thank Susanne Bauch for preparing the vimentin protein, Anna V. Schepers for providing the comparison of vimentin mechanics measurements in different buffer conditions, Monika Raabe for carrying out the phosphopeptide enrichment and Harald Herrmann for helpful discussions. This work was financially supported by the European Research Council (ERC) under the European Unions Horizon 2020 research and innovation program (Consolidator Grant Agreement no. 724932, to S.K.). Further financial support was received from the Deutsche Forschungsgemeinschaft (DFG) in the framework of SFB 860 (project number B10, to S.K.), SFB 1286 (project number A08, to H.U.) and of Germany’s Excellence Strategy - EXC 2067/1-390729940 (MBExC, to S.K. and B.S.). C.L. received a fellowship of the Studienstiftung des deutschen Volkes.

## Author Contributions

S.K. conceived and supervised the project. J.K. performed the experiments and analyzed the data. J.M. and B.S. provided the 14–3–3 protein. H.U., I. P. and I. S. performed the mass spectrometry measurements. C.L. performed the numerical simulations. M.D. performed the small-angle X-ray scattering experiments. J.K. and S.K. wrote the manuscript with contributions from all authors.

## Data availability

The data that support the findings of this study are available from the corresponding authors upon reasonable request.

