## Supplementary material for "Post-Translational Modifications Soften Vimentin Intermediate Filaments": SI

⊥*Max Planck School "Matter to Life".*

#### Theoretical Model and Monte-Carlo Simulation

To simulate the force-strain behavior of vimentin IFs, we calculate the spring constants of the modeled elements and the transition rates of possible reactions and run a Monte-Carlo simulation with a self-written Matlab code (MathWorks, Natick, Massachusetts, USA) as in

Refs. 1,2. We model each monomer consisting of three $\alpha$ helices as one spring with a spring constant $\kappa_\alpha$ and an element, which can elongate under force to an unfolded state $u$. To theoretically describe an entire filament, we connect these monomers *via* springs as they are associated in an actual filament: Two monomers are laterally connected to form a dimer, two dimers form a tetramer, and eight tetramers a ULF. To form a filament, 100 ULFs are placed in series and connected *via* springs with a spring constant $\kappa_L$, which represents the longitudinal bonds between ULFs.

From crystallography[3] and hydrogen exchange[4] experiments, we know that there are specific sites for dimers and tetramers to bind to each other. Therefore, in addition to the theoretical models presented previously,[1,2] we take the links between dimers and between tetramers as connecting springs into account as sketched in Fig. 18. The spring constant $\kappa_{bt}$ represents bonds between tetramers (marked in green in Fig. 18) and the spring constant $\kappa_{bd}$ represents bonds between dimers, *i.e.* within tetramers (marked in blue in Fig. 18).

To calculate the total force acting on the filament, we first determine the spring constant of the filament. The spring constant of the $i$th ULF in the filament depends on the number of intact $\alpha$ helices $A_j$ in this ULF with spring constant $\kappa_\alpha$. Upon loading, the $\alpha$ helices open into an unfolded state $u$, in which the unfolded $\alpha$ helix has the spring constant $\kappa_u$. We assume that a ULF consists of $N_P = 32$ parallel monomers[5] and $N_P/4 = 8$ parallel tetramers. The number of tetramers with intact (*i.e.* unfolded) $\alpha$ helices is $\lfloor A_j/4 \rfloor$, thus the number of possible interactions between these tetramers with intact $\alpha$ helices is $\lfloor A_j/4 \rfloor - 1$. Here, we assume that the interaction between the tetramers consisting of (formerly) intact $\alpha$ helices is lost when one $\alpha$ helix unfolds. Intact $\alpha$ helices in a tetramer which contains unfolded $\alpha$ helices are more likely to unfold than intact $\alpha$ helices in a tetramer containing only intact $\alpha$ helices. Thus, we assume that the next unfolding $\alpha$ helix under force is more likely to be located in a tetramer containing already unfolded $\alpha$ helices. For example, if 18 $\alpha$ helices in a ULF are intact, $\lfloor 18/4 \rfloor - 1 = 4 - 1 = 3$ connections between four intact tetramers are left. If another $\alpha$ helix unfolds, we assume that the unfolding occurs in the tetramer with only two intact $\alpha$ helices.

We describe a bond between tetramers with the spring constant $\kappa_{bt}$. Similarly, we model the dimer-dimer connection within a tetramer with the spring constant $\kappa_{bd}$. The number of intact dimer-dimer connections is the same as $\lfloor A_j/4 \rfloor$, because we assume that an unfolded $\alpha$ helix breaks the bonds between two dimers. Thus, the bonds between dimers and tetramers contribute $(\lfloor A_j/4 \rfloor - 1)\kappa_{bt} + \lfloor A_j/4 \rfloor \kappa_{bd}$ to the stiffness of the $A_j$ $\alpha$ helices. Longitudinally, *i.e.* along the filament, we assume "linkers" that connect the individual ULFs as well as the single $\alpha$ helices within one monomer. $\kappa_L$ represents the stiffness of these linkers, and the linkers and ULFs are connected in series. In case all $\alpha$ helices are unfolded, all monomers are in the unfolded state, which has a spring constant $\kappa_u$, *i.e.* the $N_P$ monomers have a total stiffness of $N_P \kappa_u$. Note that as long as there is at least one intact $\alpha$ helix present in a ULF, $\kappa_u$ does not contribute to the overall stiffness, as these unfolded monomers are longer and thus bear no force.

For the fully coupled $j$th ULF including the connection to the next ULF, we obtain the spring constant $\kappa_j$:

$$\kappa_j = \begin{cases} \left( \frac{1}{\kappa_L} + \frac{1}{N_P \kappa_u} \right)^{-1} & \text{for } \; A_j = 0 \\ \left( \frac{1}{\kappa_L} + \frac{1}{A_j \kappa_\alpha + \lfloor \frac{A_j}{4} \rfloor \kappa_{bd} + \left( \lfloor \frac{A_j}{4} \rfloor - 1 \right) \kappa_{bt}} \right)^{-1} & \text{for } A_j > 0 \end{cases} \tag{1}$$

Here, $\lfloor A_j/4 \rfloor$ is the number of links between dimers in the ULF and $\lfloor A_j/4 \rfloor - 1$ the number of links between tetramers. In case the lateral bonds between tetramers are broken, only the bonds between the dimers within a tetramer remain. Thus, $N_C$ subunits with $N_M$ monomers form a ULF, *e.g.* in the case of 32 monomers per ULF, if the subunits are tetramers, which are not coupled, we obtain $N_M = 4$ and $N_C = 8$. In contrast to the fully coupled ULF described with Eq. 1, we assume that subunits with only unfolded $N_M$ monomers contribute to the overall ULF stiffness as well, since there are no strong bonds inhibiting the subunit to slide past its original neighboring subunits. The stiffness of the $j$th ULF with $A_{j,m}$ as the number of intact $\alpha$ helices of the $m$th subunit in the $j$th ULF, the number $I_j$ of subunits with only unfolded $\alpha$ helices and the number of dimers coupled *via* $\kappa_{bd}$ within a subunit $N_d = \lfloor \sum_{m=1}^{N_C} A_{j,m}/N_M \rfloor$ is:

$$\kappa_j = \begin{cases} \left(\frac{1}{\kappa_L} + \frac{1}{N_P \kappa_u}\right)^{-1} & \text{for } \sum_{m=1}^{N_C} A_{j,m} = 0 \\ \left(\frac{1}{\kappa_L} + \frac{1}{N_M \kappa_u I_j + \sum_{m=1}^{N_C} A_{j,m}\kappa_\alpha + N_d \kappa_{bd}}\right)^{-1} & \text{for } \sum_{m=1}^{N_C} A_{j,m} > 0. \end{cases}$$

In the case of dimer sliding, neither dimers nor tetramers couple and $N_d = 0$:

$$\kappa_j = \begin{cases} \left(\frac{1}{\kappa_L} + \frac{1}{N_P \kappa_u}\right)^{-1} & \text{for } \sum_{m=1}^{N_C} A_{j,m} = 0 \\ \left(\frac{1}{\kappa_L} + \frac{1}{N_M \kappa_u I_j + \sum_{m=1}^{N_C} A_{j,m}\kappa_\alpha}\right)^{-1} & \text{for } \sum_{m=1}^{N_C} A_{j,m} > 0. \end{cases}$$

Since all ULFs are connected in series to form a filament, the stiffness of the filament $\kappa_F$ becomes $\kappa_F = 1/(\sum_{j=1}^{N_E} 1/\kappa_j)$.

To obtain the force-strain behavior as in Fig. 4c in the main text, we set the model parameters to the following values: $\kappa_\alpha = 6.5$, $\kappa_{bd} = 7$, $\kappa_{bt} = 7$, $\kappa_L = 60$, $\kappa_u = 20$, $N_E = 100$ and $N_P = 32$. In case of strong coupling in Fig. 4c in the main text, we assume that there is only one large subunit per ULF consisting of 32 monomers. For the less coupled case, we assume tetramers as subunits with $N_M = 4$ and $N_C = 8$ (light blue in Fig. 4c in the main text) and for the least coupled case, we assume dimers as subunits with $N_M = 2$ and $N_C = 16$ (blue in Fig. 4c in the main text). Further parameters are necessary to run the simulation, but they do not influence the spring constant of a ULF: The free energy difference between the unfolded and folded state $\Delta G = 2\ k_B T$, the normalized length change upon unfolding $\Delta L = 1$ and a factor to ensure in detailed balance $\theta = 0.9$.

To complete the calculation of the force-strain behavior, the extension of the $j$th ULF $\lambda_j$ is:

$$\lambda_j = \begin{cases} 0 & \text{if for all } m\text{: } A_{j,m} > 0 \\ 1 & \text{if for any } m\text{: } A_{j,m} = 0. \end{cases}$$

For a fully coupled filament, $m = 1$. The total extension of the filament then is $\lambda_{tot} = \sum_{i=1}^{N_E} \lambda_j$. Since the optical traps pull on the filament with a constant velocity $v$, the end-to-end distance $x(t)$ at time $t$ is $x(t) = vt$. The force $\phi$ on the filament becomes

$$\phi = \kappa_F(x - \lambda_{tot}) \,.$$

All closing and opening reaction rates of $\alpha$ helices are updated with the new value of $\phi$. The next reaction and time step is determined with the Gillespie algorithm. We repeat the procedure of spring constant calculation, length change, force increase and $\alpha$ helix reaction until we obtain a complete force-strain curve for the strain of interest.

### Materials and Methods

#### 14–3–3 purification

Recombinant maltose-binding protein (MBP)-tagged protein 14–3–3$\gamma$ was expressed and purified from *E. coli* strain BL21 Rosetta. For the actual measurements the MBP-tag was removed from 14–3–3. Protein expression was performed in 2 YT media (2YT mix, AppliChem GmbH, Darmstadt, Germany) and induced with 1 mM IPTG for 3 h at 30°C. Cells were harvested and lysed in MBP purification buffer (20 mM HEPES, pH 7.4, 150 mM KOAc, 5 mM $Mg(OAc)_2$, 1 mM EDTA, 1 mM DTT, 1 mM PMSF). Crude cell lysate was centrifuged at 100,000 g for 30 min at 4°C. The supernatant was incubated with 1 mL washed amylose affinity resin (New England Biolabs, Frankfurt, Germany) for 1 h at 4°C under gentle rotation. The bead slurry was transferred to gravity columns and first washed with 3 column volumes MBP purification buffer, pH 7.4, followed by one column volume 1 mM ATP (Carl

Roth, Karlsruhe, Germany) dissolved in MBP purification buffer, pH 7.4 and a final wash step in MBP purification buffer, pH 7.4. Bound MBP-tagged protein was eluted with 20 mM maltose elution buffer (20 mM HEPES, pH 7.4, 150 mM KOAc, 5 mM $Mg(OAc)_2$, 1 mM EDTA, 1 mM DTT, 20 mM D-maltose).

Eluted protein was dialyzed over night in MBP purification buffer, pH 7.4. The MBP-tag was cleaved from the purified MBP-14–3–3 protein with FactorXa enzyme for 16 h at 25°C according to the supplier's recommendation (Merck MilliPore, Merck KGaA, Darmstadt, Germany). The cleaved protein was separated by size exclusion chromatography on an Äkta purifier (GE Healthcare, Freiburg, Germany) and a Seperdex75 size exclusion column in MBP purification buffer, pH 7.4. Finally, protein concentrations were measured with a Bradford assay and the purified proteins were aliquoted and stored at -80°C.

#### Vimentin filament assembly

To prepare the protein for filament assembly, 200 $\mu$L unlabeled vimentin at a concentration of 1.3 mg/mL was mixed with 25 $\mu$L ATTO647N-labeled vimentin at a concentration of 0.4 mg/mL resulting in a labeling ratio of 4 %, and dialyzed from storage buffer (8 M urea, 5 mM Tris-HCl, 1 mM EDTA, 0.1 mM EGTA, 0.01 mM MAC and 250 mM KCl, pH 7.5) to 6 M urea, 5 mM Tris-HCl, pH 8.4 and then in a stepwise manner (4 M, 2 M, 1 M, 0 M urea, 30 min for each step) to 5 mM Tris-HCl, pH 8.4 with an subsequent dialysis step in fresh 5 mM Tris-HCl buffer over night. Afterwards the protein concentration was adjusted to about 0.2 mg/mL. To initiate filament assembly, the protein was dialyzed into assembly buffer containing 25 mM Tris-HCl, pH 7.5 and 50 mM NaCl at 37°C over night.[6]

#### Phosphorylation of vimentin

As phosphorylation buffer we used 25 mM Tris-HCl, pH 7.5 containing 50 mM NaCl, 2 mM $MgCl_2$ and added 100 $\mu$M ATP (Carl Roth), and protein kinase A (PKA; New England Biolabs). The amount of PKA was dependent on the amount of vimentin used, with 1 $\mu$L

PKA solution per 1 $\mu$g vimentin. Vimentin solution and phosphorylation buffer were mixed at a ratio of 1:9 resulting in a final vimentin concentration of 0.11 mg/mL. This mixture was incubated for about 1 h at 37°C and dialyzed into 8 M urea in 5 mM Tris-HCl, pH 8.4 for about 1 h at room temperature. This step was performed to stop the phosphorylation by inactivation of the PKA. In a next step the phosphorylated vimentin was mixed at the desired ratios with unphosphorylated vimentin, which was diluted to the same concentration by adding 8 M urea in 5 mM Tris-HCl, pH 8.4. This vimentin mixture was then dialyzed as described before from 8 M urea in 5 mM Tris-HCl, pH 8.4 in steps of 4 M, 2 M, 1 M and 0 M urea to 5 mM Tris-HCl, pH 8.4 and afterwards assembled in 25 mM Tris-HCl, pH 7.5 and 50 mM NaCl at 37°C over night.

#### Determination of the degree of phosphorylation in tetramers and filaments

To test whether the phosphorylation of vimentin tetramers was successful, phosphorylation analysis gels (Phos-tag Acrylamide AAL-107, FUJIFILM Wako Chemicals Europe GmbH, Neuss, Germany) were used. These gels show additional bands above the actual protein band if the protein is phosphorylated. The phosphorylation analysis SDS gels were produced according to the manufacturer's instructions. Samples were prepared by mixing 13 $\mu$L protein with 7 $\mu$L sample buffer (150 $\mu$L Laemmli SDS sample buffer (Alfa Aesar, Kandel, Germany), 60 $\mu$L 1 M DTT), followed by an incubation at 95°C for 5 min. The gel was loaded with 15 $\mu$L of each sample and run at a constant current of 40 mA for about 35 min. Afterwards the gel was stained (InstantBlue, Sigma-Aldrich, Munich, Germany) for 1 h followed by several washing steps with water.

To check the incorporation of the phosphorylated monomers within the filament, ultracentrifugation was performed. The partially phosphorylated vimentin filaments were centrifuged at 34,000 rpm for 10 min (rotor: Fiberlite F50L-25x1.5; centrifuge: Sorvall WX80+ Ultra Series centrifuge, Thermo Fisher Scientific, Kandel, Germany). The supernatant was removed and mixed with sample buffer as described above. The pellet was dissolved in 8 M urea, 5 mM Tris-HCl, pH 8.4, with subsequent dialysis (8 M, 4 M, 2 M, 1 M, 0 M urea in 5 mM Tris-HCl, pH 8.4). Afterwards the dissolved pellet was mixed with sample buffer followed by an incubation at 95°C for 5 min for all samples. The samples were then loaded on an SDS gel and it was run and stained as described above.

**Binding 14–3–3 to vimentin filaments** To bind the protein 14–3–3 to vimentin it was first diluted to the same concentration determined in g/L as the vimentin solution. Then, 14–3–3 and assembled vimentin filaments were mixed at a ratio of 1:1 with respect to the concentration in g/L and incubated for 1 h at 37°C. For the optical trap measurements, 30 $\mu$L of this solution were diluted by 1 mL assembly buffer (25 mM Tris-HCl, pH 7.5 and 50 mM NaCl).

#### Verification of vimentin binding to 14–3–3

To test whether the binding of 14–3–3 to vimentin was successful, a streptavidin pulldown assay was performed (adapted from Ref. 7). Unless otherwise stated, a centrifugation speed of $200 \times$ g for 30 s was used. First 200 $\mu$L biotin-labeled vimentin (labeling with biotin-maleimide (Jena BioSciences GmbH, Jena, Germany) was mixed according to the protocol described in Ref. 8), dialyzed to tetramers and phosphorylated as described above. This biotin labeling of vimentin is necessary for the binding to the beads. The streptavidin-agarose beads (Sigma Aldrich) were washed. To do so, 70 $\mu$L beads (for 1 $\mu$g of vimentin) were pipetted into a reaction tube (1.5 mL) and washed three times with phosphorylation buffer (25 mM Tris-HCl, pH 7.5 containing 50 mM NaCl, 2 mM $MgCl_2$). In-between the washing steps the beads were centrifuged down for 30 s at 1,700 rpm (MiniSpin F-45-12-11, Eppendorf, Wesseling-Berzdorf, Germany) and the supernatant was discarded. Vimentin was diluted with phosphorylation buffer to a concentration of 0.02 mg/mL and then 50 $\mu$L of vimentin solution (total protein amount of 1 $\mu$g) was mixed with the beads in the reaction tube. The solution was incubated for 1 h at 8°C on a rotation wheel. During this time, vimentin bound to the beads due to the biotin-streptavidin binding. In order to remove the unbound vimentin, this mixture was pipetted on a column (35 $\mu$m pore size, MoBiTec GmbH, Göttingen, Germany), centrifuged down and the flow-through was kept for later analysis. The bead mixture was then again washed twice with phosphorylation buffer and the flow-through was kept. Afterwards, 14–3–3 was diluted to 0.02 mg/mL in phosphorylation buffer containing 0.01 % Triton X-100 and 50 $\mu$L were mixed with the beads to which vimentin was bound. This mixture of beads and 14–3–3 was incubated for 1 h at 8°C on a rotation wheel so the 14–3–3 bound to the vimentin on the beads. To remove the unbound 14–3–3, the mixture was centrifuged down, and the flow through was kept for later analysis. The bead mixture was then washed twice with phosphorylation buffer. To remove the bound vimentin and 14–3–3, eluation buffer was used. The eluation buffer consisted of 90 $\mu$L SDS loading buffer, 10 $\mu$L fresh DTT and 5 $\mu$L 100 mM biotin which was mixed and incubated for 10-15 min at 95°C. This eluation buffer was added to the beads, mixed well and incubated for 7 min at 70°C. In the next step the mixture was centrifuged down and the flow through was kept as it should contain the vimentin which bound to the beads and the 14–3–3 which bound to the vimentin. As a last step all the flow through samples were mixed with sample buffer and a phosphorylation analysis SDS gel was run as described above.

#### Data sets

In total, 43 control measurements with untreated vimentin were performed. For the filaments containing 1 % phosphorylated monomers 38 measurements were performed, for the ones with 5 % phophorylated monomers 41 measurements were performed and for the ones with 10 % phosphorylated monomers 38 measurements were performed. Measurements with the filaments containing 5 % phosphorylated monomers and incubated with 14–3–3 were performed 33 times.

For the mutant S38E, 34 measurements were performed with 1 % of the mutant, 33 measurements with 5 % of the mutant, 30 measurements with 10 % of the mutant and 24 measurements with 5 % mutant incubated with 14–3–3.

For the mutant S72E, 30 measurements were performed with 1 % of the mutant, 30 measurements with 5 % of the mutant, 32 measurements with 10 % of the mutant and 28 measurements with 5 % mutant incubated with 14–3–3.

#### Mass spectrometry

##### Cross-linking Experiments

Phosphorylated and non-phosphorylated vimentin was cross-linked in presence of 14–3–3 to determine the interaction sites of the two proteins after phosphorylation. First, the optimal cross-linker–to–protein ratio was determined by using 2.6 $\mu$g/2.4 $\mu$M aliquots of the complex and the individual proteins and a molar excess of the cross-linker ranging from 20 to 1,000-fold as well as a non-cross-linked control. The cross-linking reaction was performed with freshly prepared bis(sulfosuccinimidyl)suberate ($BS^3$, 100 mM stock in DMSO, Thermo Fisher Scientific) for 30 min at room temperature. The reaction was quenched by addition of Laemmli sample buffer and samples were analyzed by sodium dodecyl sulfate polyacrylamide gel electrophoresis (SDS-PAGE) on a 4–12% gradient gel (Invitrogen, Kandel, Germany). After coomassie staining, unique shifted bands were observed for a complex of phosphorylated vimentin and 14–3–3 corresponding to a heterodimer and -tetramer as in Fig. 17. For the main experiment, samples were cross-linked with a 500- and 1,000-fold molar excess of $BS^3$ and the shifted bands mentioned above were cut, in-gel digested, and peptides were extracted as described elsewhere.[9]

A quantitative cross-linking approach was pursued to examine the structural changes of vimentin caused by phosphorylation. Therefore, phosphorylated, and non-phosphorylated vimentin samples were cross-linked with differentially isotope-labeled disuccinimidyl suberate (DSS) containing either zero or four deuterium atoms. After 30 min at room temperature, the reaction was quenched with 50 mM Tris, pH 8.1, for 15 min. Phosphorylated and non-phosphorylated vimentin samples cross-linked with the opposite isotopic labels were mixed in equal ratios and the labels were swapped for a second reaction replicate. Subsequently, proteins were precipitated with chloroform and methanol[10] and resuspended in 8 M urea. After complete resuspension, samples were diluted to 4 M urea with 100 mM ammonium bicarbonate and reduced and alkylated with 10 mM dithiothreitol and 55 mM iodoacetamide in 50 mM ammonium bicarbonate, respectively. Samples were diluted to 1 M urea and digested with trypsin overnight in a 1:20 (w/w) ratio. Peptides were desalted with C18 micro spin columns (Harvard Apparatus, Holliston, Massachusetts, USA) and dried in a vacuum centrifuge (Savant SpeedVac Concentrator, Thermo Fisher Scientific).

##### Phosphopeptide enrichment

Phosphorylated vimentin sample was reduced and alkylated with 10 mM dithiothreitol and 55 mM iodoacetamide in 50 mM ammonium bicarbonate. The sample was digested overnight using trypsin at a trypsin-to-protein ratio of 1:20 (w/w) and then concentrated in the vacuum centrifuge. An aliquot of the sample was subjected directly to LC-MS/MS analysis. For phosphopeptide enrichment, TiO2-beads (10 $\mu$m, GL Science) were resuspended in buffer A: 80 % acetonitrile (v/v) 5 % trifluoroacetic acid (TFA, v/v/) 5 % Glycerol (v/v) in water. The bead suspension was mounted onto a plastic pipette tip with a filter paper support forming an approximately 3 mm long chromatographic column. The beads were equilibrated with buffer B (80 % acetonitrile (v/v) 5 % TFA (v/v) in water and 60 %) and buffer A sequentially. The sample was dissolved in 60 $\mu$L buffer A and applied onto the column. Next, the column was washed three times with buffer A and buffer B, followed by a wash with buffer B2 (60 % acetonitrile (v/v), 0.1 % TFA (v/v) in water). The retained phosphopeptides were eluted using 0.3 N NH4OH in water (pH 10.5). The sample was acidified using 10 % TFA (v/v) in water and dried in a vacuum centrifuge.

**LC-MS analysis**

Dried peptides were dissolved in 5 % (v/v) acetonitrile, 0.1 % (v/v) trifluoroacetic acid, and injected in technical duplicate (cross-linked sample) or as a single injection (unmodified and phosphorylated peptides) onto a C18 PepMap100 $\mu$-Precolumn (0.3 x 5 mm, 5 $\mu$m, Thermo Fisher Scientific) connected to an in-house packed C18 analytical column (75 $\mu$m x 300 mm; Reprosil-Pur 120C18-AQ, 1.9 $\mu$m, Dr. Maisch GmbH, Ammerbuch, Germany). Liquid chromatography was operated on an UltiMate 3,000 RSLC nanosystem (Thermo Fisher Scientific). For the cross-linked sample, a linear gradient of 10 to 50 % buffer B (80 % (v/v) acetonitrile, 0.08 % (v/v) formic acid) was applied at 300 nL/min flow rate, and 43 min total gradient duration. Eluting peptides were sprayed into a QExactive HF-X (Thermo Fisher Scientific) mass spectrometer. MS1 scans were performed with a scan range from $m/z$ 350 to 1,600, a resolution of 120,000 full width at half maximum (FWHM), $1 \times 10^6$ automatic gain control (AGC) target, and 50 ms maximum injection time. Each MS1 scan was followed by 20 MS2 scans of the most abundant precursors fragmented with a normalized collision energy of 30 and acquired with a resolution of 30,000 (FWHM), $1 \times 10^5$ AGC target, and 128 ms maximum injection time. Only charge states from 3+ to 8+ were considered, and a dynamic exclusion of 20 s was set. Vimentin cross-linked with isotopically labeled DSS was analyzed identically with the exception of 30 s dynamic exclusion time.

Similarly, vimentin peptides before and after the titanium dioxide enrichment step were analyzed using a 73 min long linear gradient from 10 to 42 % of the buffer B. The samples were sprayed into a QExactive (Thermo Fisher Scientific) mass spectrometer operated at 70,000 resolution, $1 \times 10^6$ AGC target, and 50 ms maximum injection time for MS1 scans; and 17,500 resolution, $1 \times 10^5$ AGC target and 54 ms maximum injection time for MS2 scans. Per MS1 scan, 20 peptide precursors with charge states 2-6 were selected for fragmentation using normalized collision energy of 30 %. An isolated precursor ion was excluded from repetitive selection for 25 s.

#### Data analysis for mass spectrometry

Raw files were submitted to a cross-link database search with pLink 2 (version 2.3.9)[11] against the sequences of human vimentin and 14–3–3 protein $\gamma$. The following search parameters were defined: maximum three missed cleavages, cysteine carbamidomethylation as fixed modification, methionine oxidation and phosphorylation of serine, threonine and tyrosine as variable modifications, 4 to 100 amino acids peptide length, 400 to 10,000 Da peptide mass, 10 ppm and 20 ppm precursor and fragment ion mass deviation, respectively, 10 ppm filter tolerance, 1 % false discovery rate cut-off and a cross-linker reactivity towards lysine, serine, threonine and tyrosine. Database search results were filtered for at least 4 matched fragment ions per peptide in a pair and a minimum score of 1 (negative decadic logarithm of the initial score).

Quantitative cross-linking acquisitions were analyzed with pLink1 (version 1.23)[12] after a conversion to mgf format with Proteome Discoverer version 2.1 (Thermo Fisher Scientific). Data were searched with the same parameters except the following: a 25 ppm precursor ion and 10 ppm filter tolerance window around the monoisotopic and the first, second and third isotopic mass, no phosphorylation as variable modification, 1 % false discovery rate cut-off and no further filtering post-search. Quantification was performed with XiQ[13] by extracting areas under the curve of the first to third isotopic peak of an identification demarcated by a decrease to 10 % signal intensity. Abundance ratios were $\log_2$-transformed and median normalized. The leading sign of ratios was changed for the label-swap replicate. Quantified redundant cross-link-to-spectrum matches were then merged to unique cross-linked residues with a custom R script as described previously.[14] Briefly, median ratios were calculated for each charge state per peptide, which were then summarized to unique peptides as a weighted average. Unique peptides were finally summarized to unique linked residues as median ratios of all supporting peptides. Cross-links were visualized on proteins with xiNET[15] and quantitative values were plotted with Perseus.[16]

Analysis of phosphorylated and non-phosphorylated peptides was performed in MaxQuant version 1.6.2.10[17,18] using reviewed human protein sequences from Uniprot (02/2019)[19] supplemented with the modified vimentin sequence. Cysteine carbamidomethylation was set as fixed modification; methionine oxidation, protein N-term acetylation and phosphorylation of serine, threonine and tyrosine were selected as variable modifications. Maximum of two missed cleavage sites and up to five variable modification were allowed per peptide. Other settings were kept default. Peptide intensities were extracted as area under the chromatographic peak using Skyline version 19.1.0.193.[20] Intensities of phosphorylated peptides were normalized by intensities of the respected non-phosphorylated peptides using a custom R script.

#### SAXS experiments

Assembly for SAXS experiments was performed in 1.5 mm diameter quartz glass capillaries, wall thickness 0.01 mm (Hilgenberg GmbH, Malsfeld, Germany) for 4 h at 37 °C in a temperature controlled water bath, because assembled protein is too viscous to be inserted properly into the capillaries. Directly after filling of the capillaries, they were sealed with wax (Hampton Research, Aliso Viejo, CA, USA).

##### SAXS measurements and data treatment

SAXS measurements were performed using an in-house SAXS setup (Xeuss 2.0, Xenocs, Sassenage, France) equipped with a Genix 3D source (Xenocs), at a wavelength of 1.54 Å (Cu $K_\alpha$ radiation), 50 kV and 600 $\mu$A. The beam was focused down to 500 x 500 $\mu m^2$. At a sample to detector distance of 1225 mm the scattered signal was collected on a Pilatus3 R 1M pixel detector (981 x 1043 pixels, pixel size 172 x 172 $\mu m^2$ Dectris Ltd., Baden, Switzerland). To block the primary beam, a 3 mm-diameter beamstop was placed directly in front of the detector. In total, the signal was recorded for 12 h, divided into 15 min intervals. To obtain the background signal, which is needed for a proper background subtraction, the buffer was measured prior to the protein in the same capillary. As a first step in data analysis, the

2D detector images were azimuthaly integrated and the background was subtracted using self-written Matlab scripts (Matlab2017a, The MathWorks, Natick, MA, USA) based on the cSAXS Matlab base package, available at https://www.psi.ch/en/sls/csaxs/software. The data were normalized to the thickness of the capillary, the exposure time, the transmission values, the correction factor and the protein concentration. The integrated intensity $I(q)$ is plotted against the magnitude of the scattering vector $q$:

$$q = \frac{4\pi}{\lambda} sin(\theta), \tag{2}$$

where $\lambda$ is the wavelength of the radiation and $2\theta$ is the scattering angle. SAXS data are shown in a range from $0.08\,\mathrm{nm}^{-1}$ to $2.00\,\mathrm{nm}^{-1}$, corresponding to real space length scales from 3.14 nm to 78 nm. We analyzed the data by performing a Guinier analysis using the software package PRIMUS[21] (ATSAS, EMBL, Hamburg, Germany) as well as the open source Python library alea.[22] Performing the Guinier analysis, the radius of gyration of the cross-section $R_c$ as well as the forward scattering $I(0)$ can be retrieved using

$$I(q) = \frac{L\pi}{q} I(0) \exp\left(-\frac{q^2 R_c^2}{2}\right), \tag{3}$$

with $L$ as the length of the object. For analysis, the small $q$-values are fitted up to a limit of $qR_g \leq 1.3$.[23,24] $R_c$ can be interpreted as the average distance from the center of gravity of the particle. In a simplified picture, vimentin filaments can be described as cylinders, for which the relation $R_c = \frac{1}{\sqrt{2}} R$ is valid. However, vimentin filaments have been shown to be partially hollow cylinders.[25,26] Taking this into account, $R_c = \sqrt{\frac{1}{2}(R_i^2 + R^2)}$ with $R_i$ and $R$ as the inner and outer radii of the cylinder, respectively. This brings the calculated values for the radius of gyration of the cross-section $R_c$ closer to the real radius of the filament $R$. In addition, it has been reported that the tails of vimentin protrude from the filament itself,[27–29] which leads to an increase of $R_c$. Therefore, we assume that $R_c$ and $R$ are very similar and thus use $R_c$ from the Guinier analysis to calculate the Young's modulus.

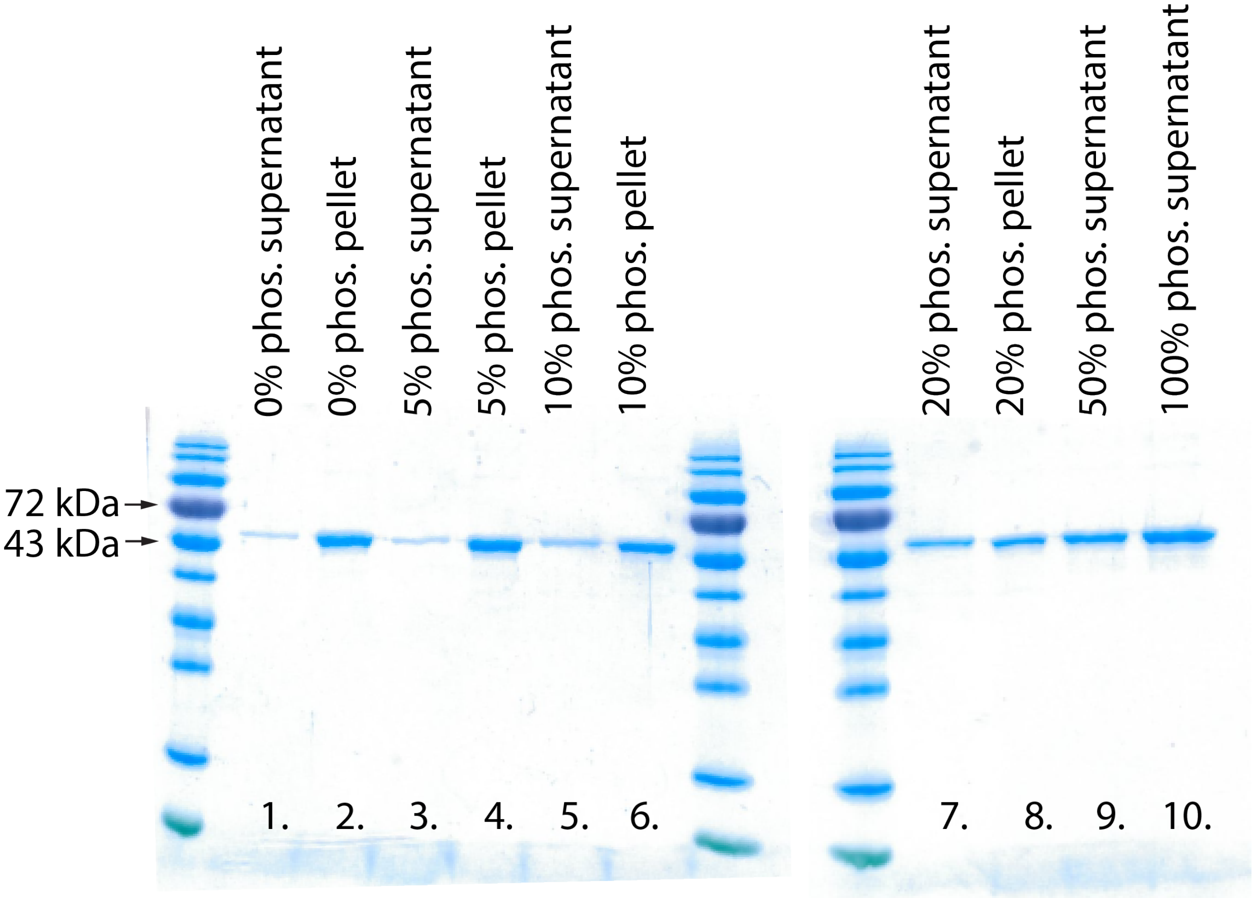


Figure 1: Determination of the maximum phosphorylation ratio that still allows for filament assembly. SDS gel of vimentin with different ratios of phosphorylated monomers ranging from 0 % to 100 %. Lane 1 shows the supernatant of vimentin without phosphorylation, lane 2 shows the pellet of the same protein after ultracentrifugation. Lane 3-4 show the corresponding data for vimentin mixture of 5 % phosphorylation, lane 5-6 for 10 % phosphorylation, and lane 7-8 for 20 % phosphorylation. Lane 9 and 10 show the supernatant of vimentin solutions with 50 % and 100 % phosphorylation. The data show that from 20 % phosphorylation on, the amount of protein in the supernatant strongly increases. Therefore, a maximum percentage of 10 % phorphorylated protein was chosen for the experiments described in the main text. It should be noted that the given percentage of phosphorylation refers to the amount of phosphorylated vimentin mixed with unphosphorylated vimentin and the actual percentage is lower as fully phosphorylation can not be achieved.

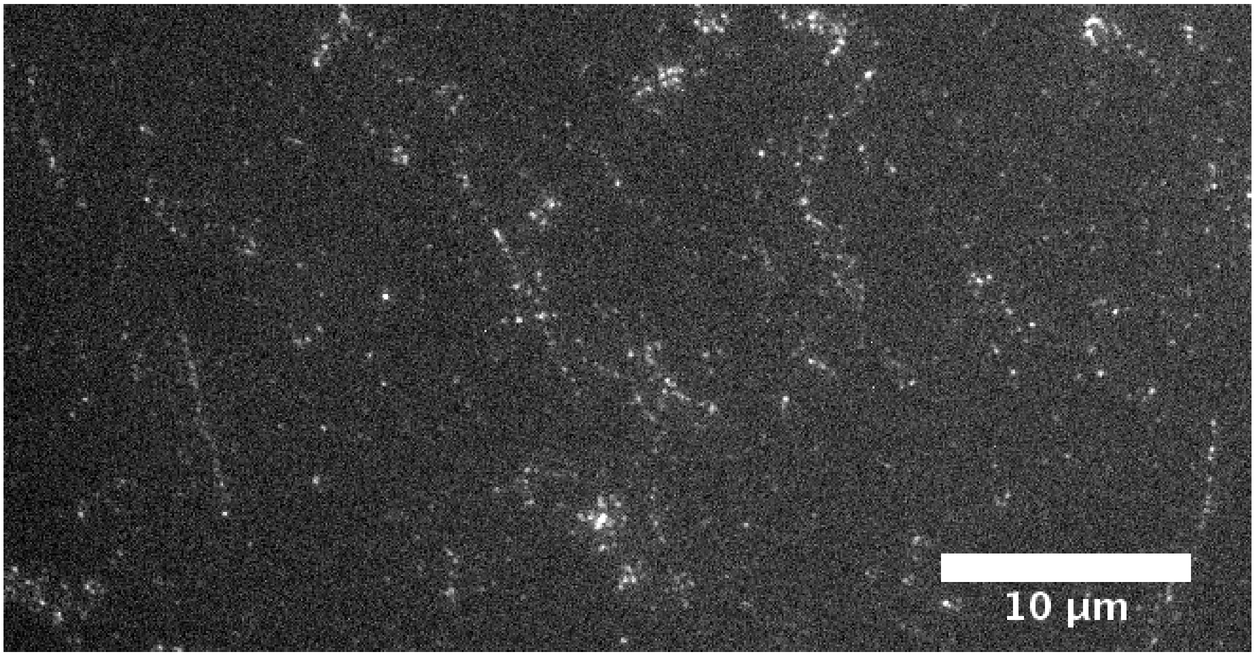


Figure 2: Incorporation of phosphorylated vimentin into filaments. Labeled vimentin is phosphorylated and mixed with unlabeled and unphosphorylated vimentin monomers at 5 %, and assembled into filaments. Dotted, but fairly long vimentin filaments can be seen and this ensures that phosphorylated vimentin monomers are incorporated into assembled filaments.

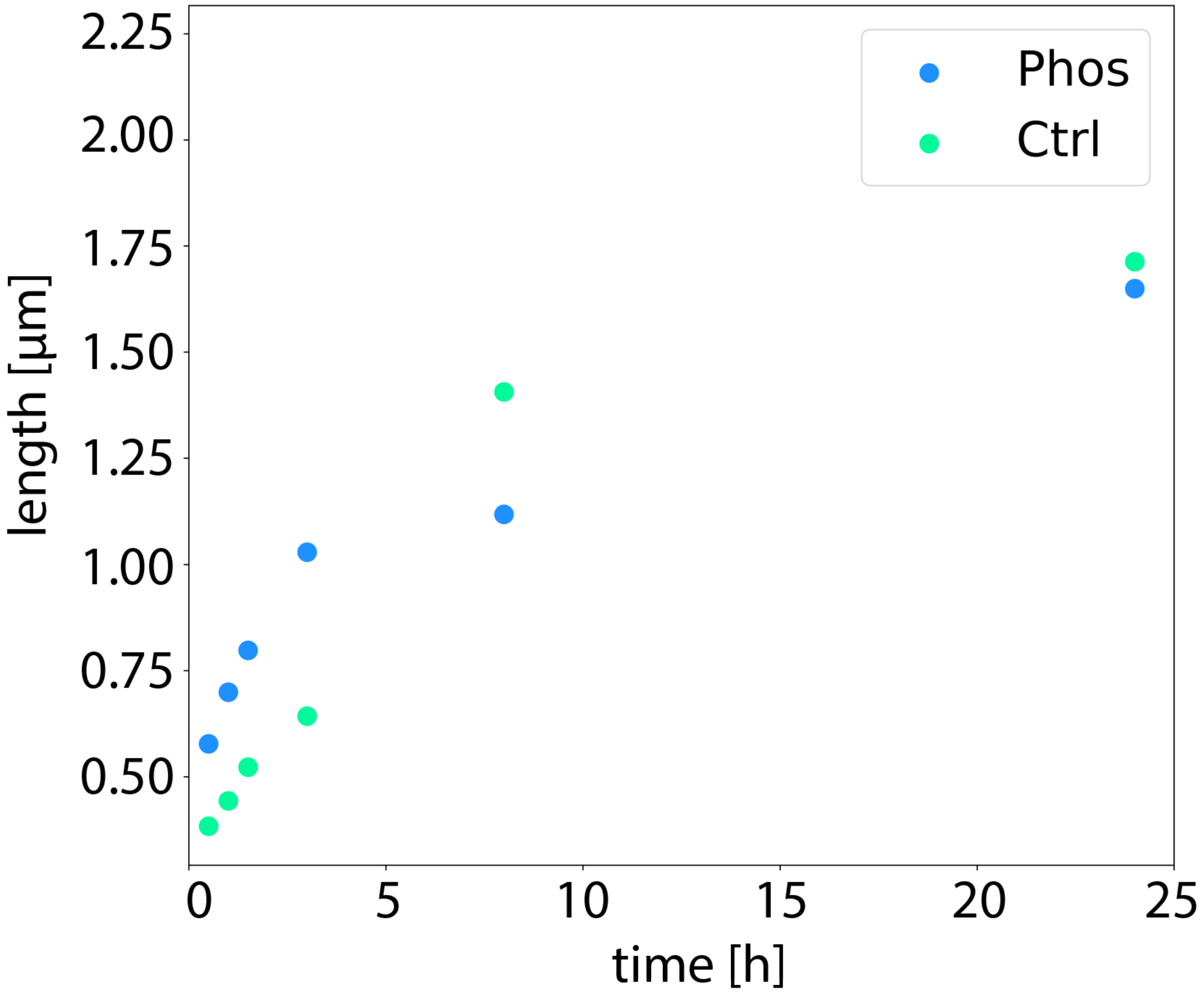


Figure 3: Length distribution of the filaments during the assembly. Untreated filaments are shown in green and filaments with 10% of phosphorylation are shown in blue. After the full assembly time of 24 h at room temperature, the average length obtained for both conditions is very similar.

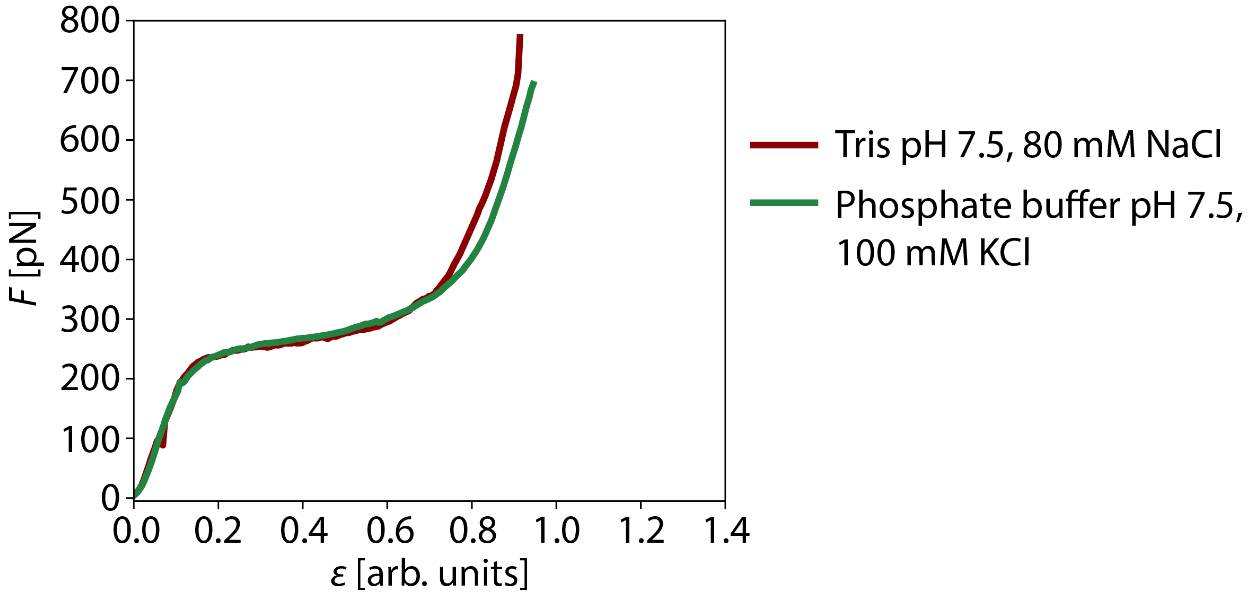


Figure 4: Comparison of force-strain curves for vimentin in different buffers. Mean curves for both conditions are shown. In dark red, vimentin force-strain curves in 25 mM Tris buffer containing 80 mM NaCl are shown. In green, vimentin force-strain curves in 2 mM phosphate buffer containing 100 mM KCl are shown. There is no influence of the buffers on the stretching behavior.

(4) Premchandar, A.; Mücke, N.; Poznański, J.; Wedig, T.; Kaus-Drobek, M.; Her-

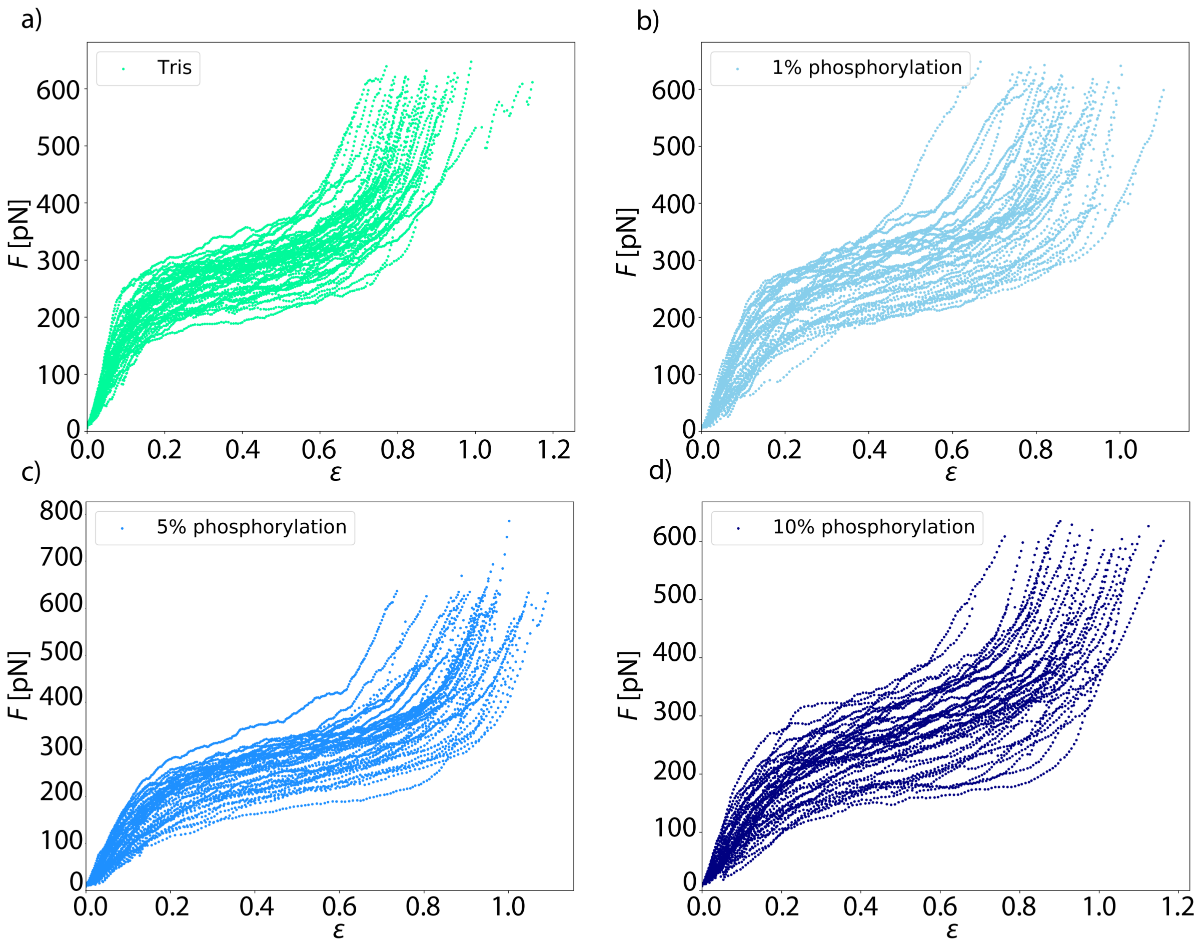


Figure 5: Single force-strain curves of all phosphorylation data. The control measurement is shown in green, filaments with 1 % phosphorylation are shown in light blue, filaments with 5 % phosphorylation are shown in medium blue and filaments with 10 % phosphorylation are shown in dark blue.

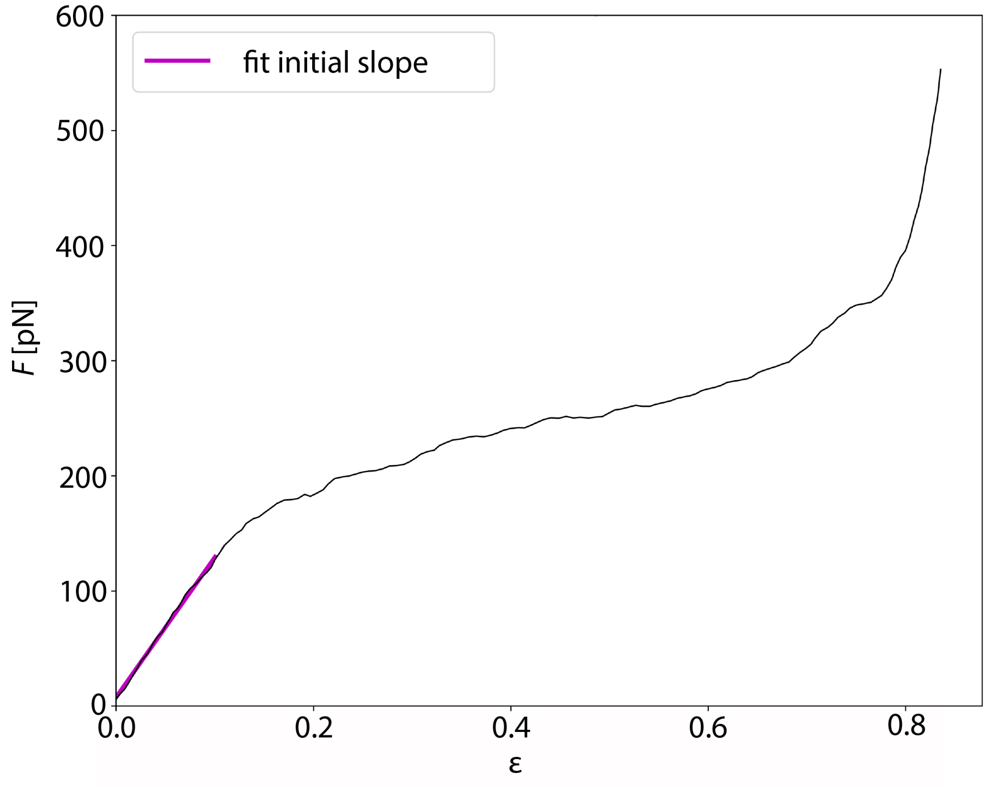


Figure 6: Determination of the initial slope used to calculate the Young's modulus. In black, a typical force-strain curve of vimentin is shown. In purple, the fit of the initial slope up to a force of 130 pN is shown.

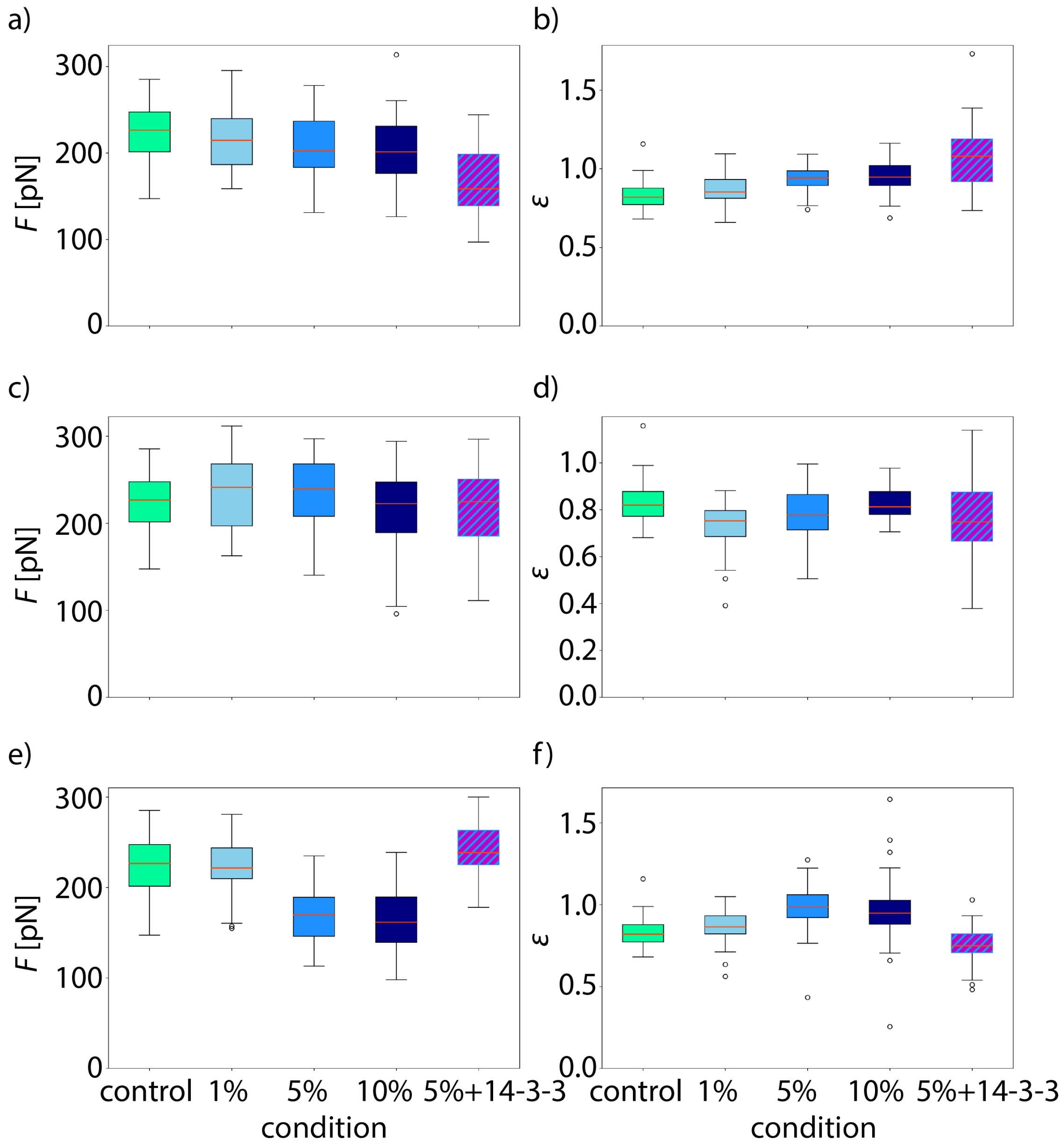


Figure 7: Additional analyses of the force-strain curves. a)-b) Vimentin filaments with different percentages of phosphorylation. a) The force $F$ at the start of the plateau. With increasing phosphorylation, the plateau starts at lower forces. b) The maximum strain $\varepsilon$ increases with increasing amount of phosphorylation. c)-d) Vimentin filaments with different percentages of mutation S38E. c) The force at the onset of the plateau stays rather constant for the different conditions. d) The maximum strain does not change for the different conditions. e)-f) Vimentin filaments with different percentages of mutation S72E. e) The force at the beginning of the plateau increases with increasing amount of the mutation S72E but increases again when filaments are incubated with 14–3–3. f) The maximum strain increases with increasing amount of the mutation S72E but decreases again when filaments are incubated with 14–3–3.

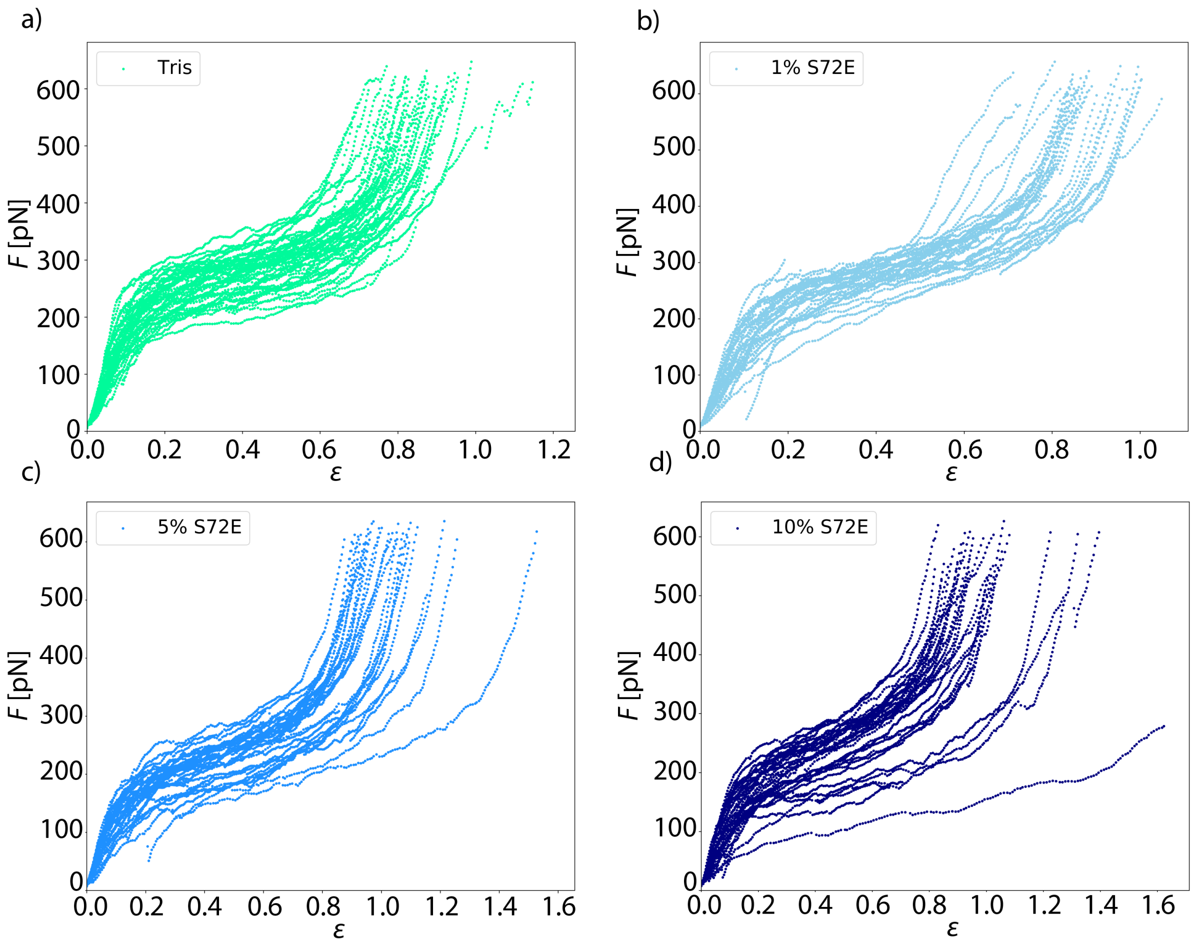


Figure 8: Single force-strain curves of all the S72E mutant data. The control measurement is shown in green, filaments with 1 % S72E mutation are shown in light blue, filaments with 5 % S72E mutation are shown in medium blue and filaments with 10 % S72E mutation are shown in dark blue.

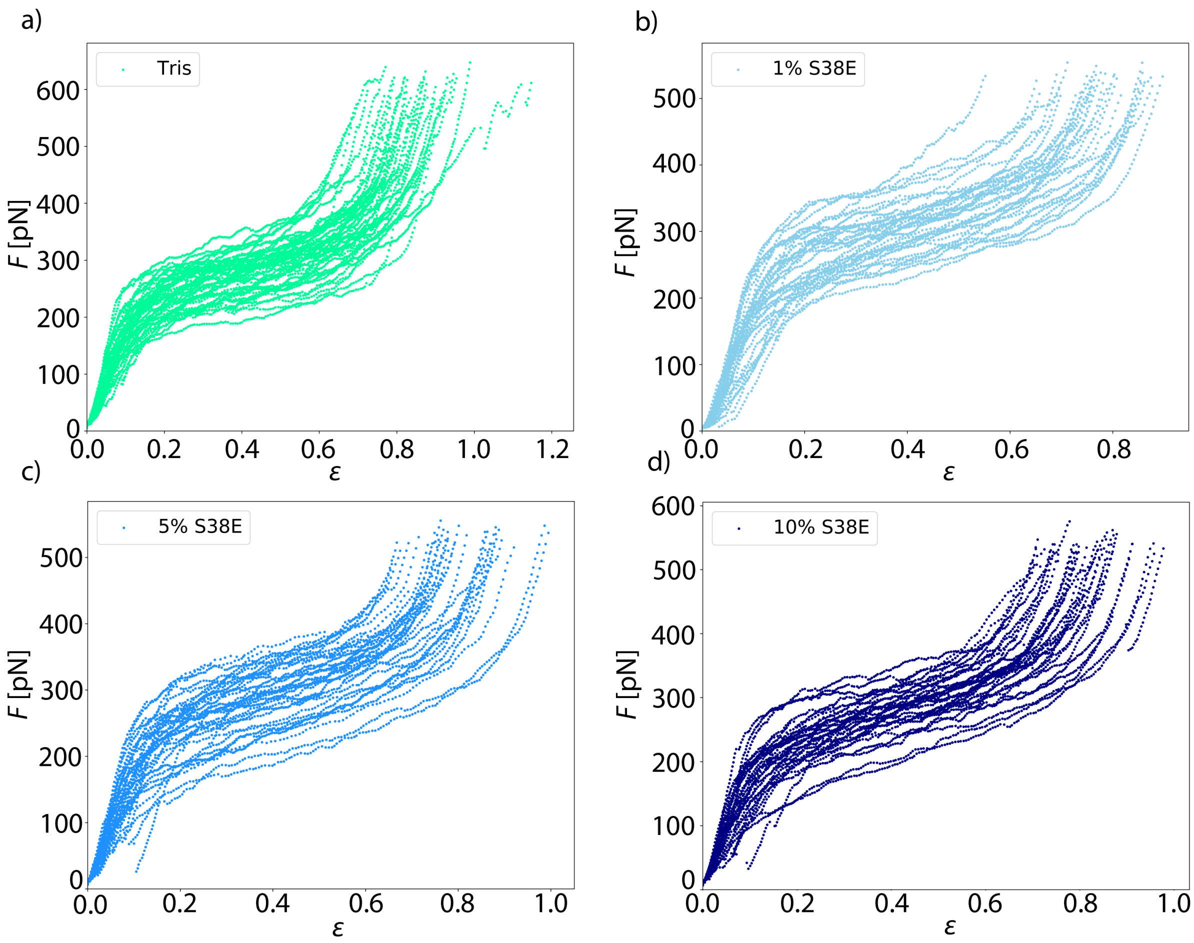


Figure 9: Single force-strain curves of all the S38E mutant data. The control measurement is shown in green, filaments with 1 % S38E mutation are shown in light blue, filaments with 5 % S38E mutation are shown in medium blue and filaments with 10 % S38E mutation are shown in dark blue.

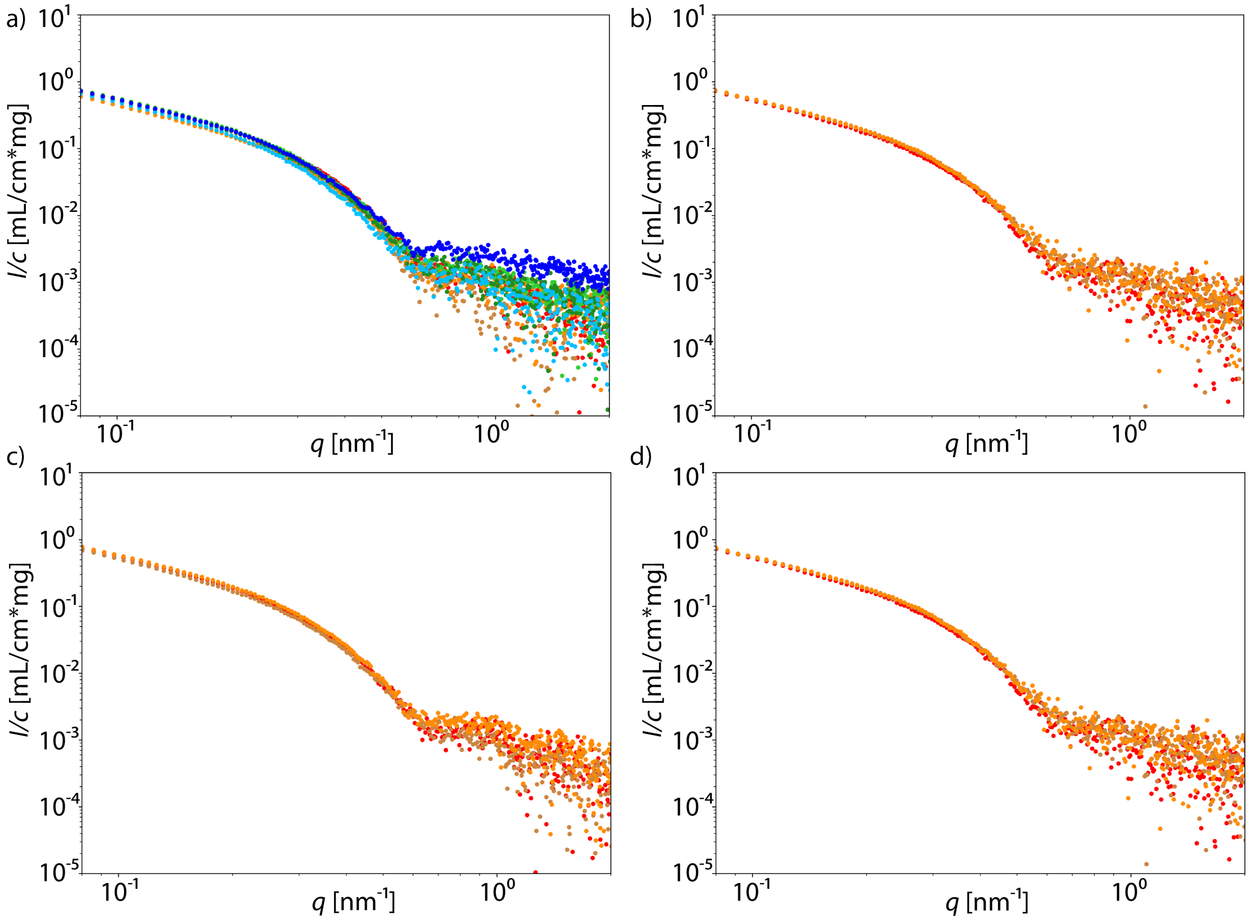


Figure 10: Individual SAXS curves of phosphorylated vimentin. Different colors denote individual experiments; a) control measurements with unphosphorylated vimentin; b) 1 % phosphorylated vimentin; c) 5 % phosphorylated vimentin; d) 10 % phosphorylated vimentin.

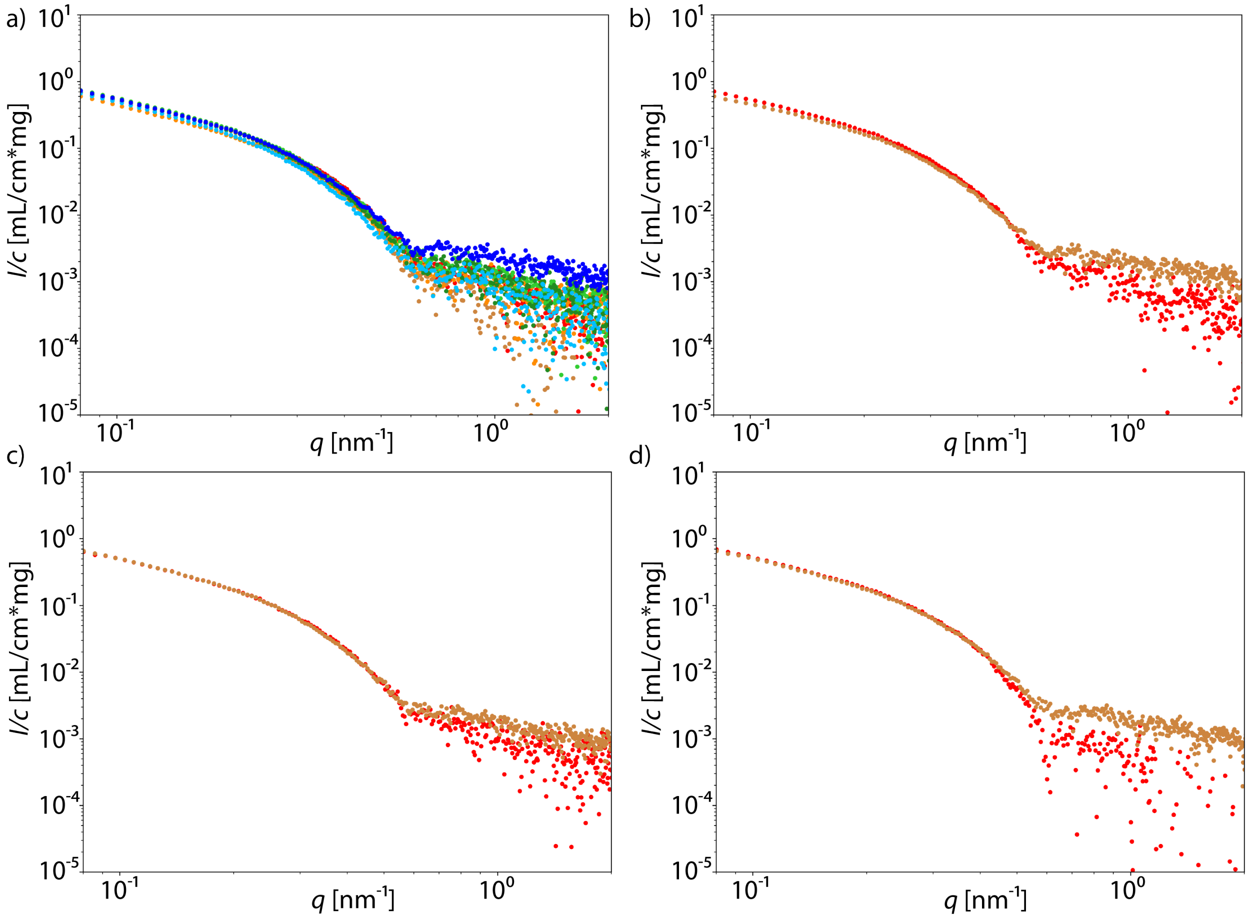


Figure 11: Individual SAXS curves of phophomimetic vimentin S38E. Different colors denote individual experiments; a) control measurements with wildtype vimentin; b) 1 % vimentin S38E; c) 5 % vimentin S38E; d) 10 % vimentin S38E.

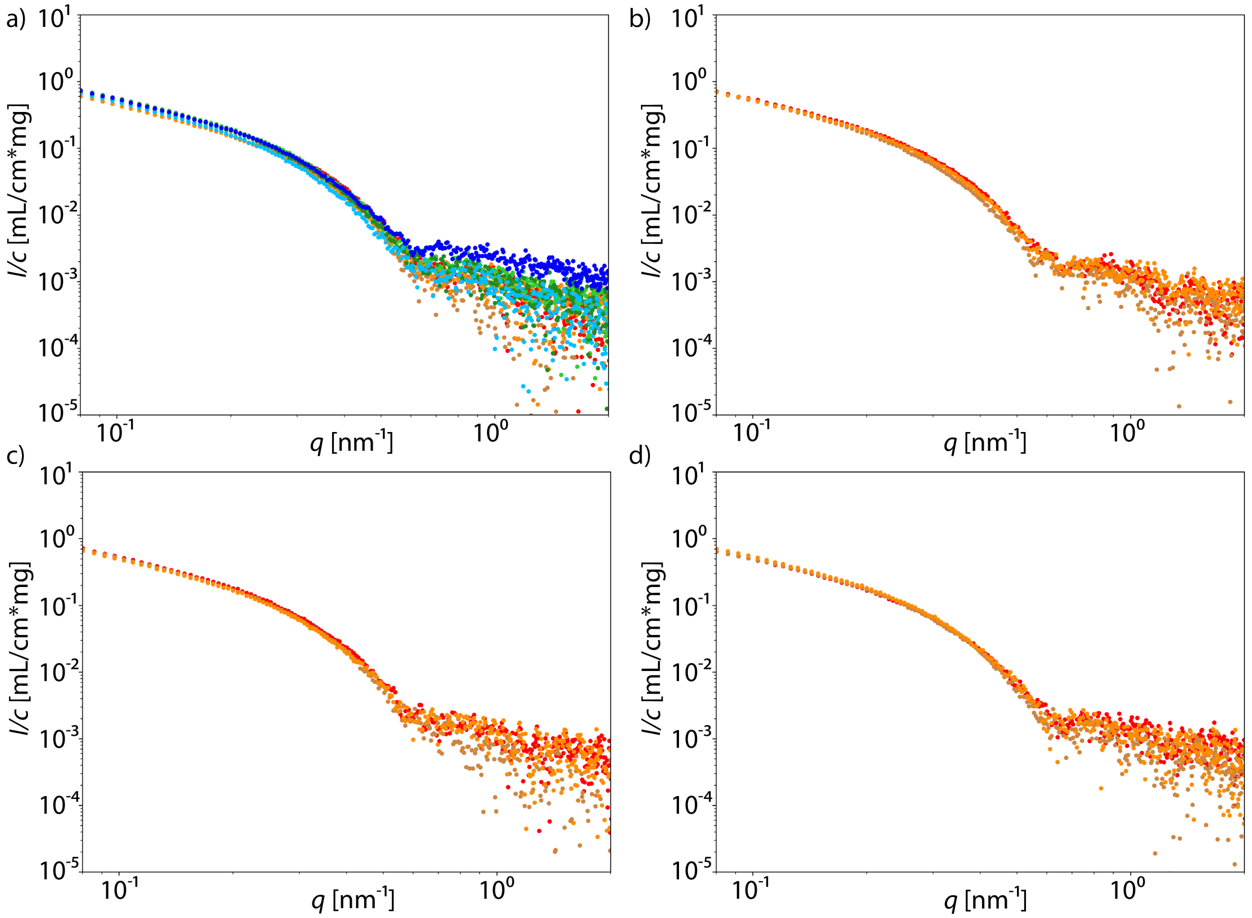


Figure 12: Individual SAXS curves of phophomimetic vimentin S72E. Different colors denote individual experiments; a)control measurements with wildtype vimentin; b)1 % vimentin S72E; c) 5 % vimentin S72E; d) 10 % vimentin S72E.

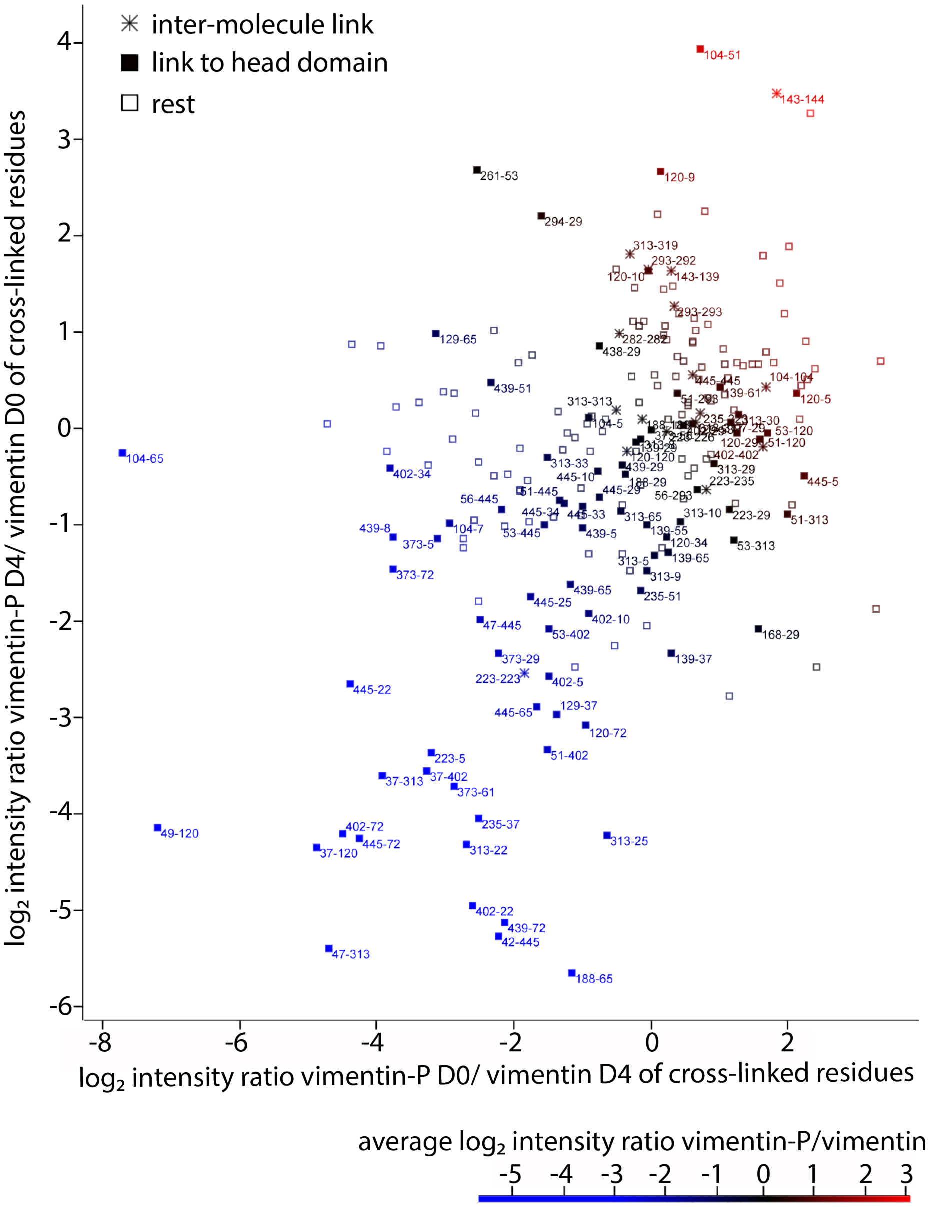


Figure 13: Quantified ratios (phosphorylated to non-phosphorylated) of cross-linked vimentin residues in two independent reaction replicates. Data point labels specify the cross-linked amino acid residues within vimentin. Cross-links including the N-terminal head domain (residue 1 to 94) are shown as filled squares and inter-molecular cross-links to the same amino acid residue are shown as asterisks. Data points are color-coded according to the average $\log_2$ intensity ratio ranging from a decreased (blue), unaltered (black), to increased (red) cross-link abundance upon phosphorylation. The majority of cross-linked residues does not change their abundance in response to phosphorylation. However, those cross-linked residues that change their abundance are mostly less abundant upon phosphorylation and are almost exclusively cross-links to the head domain. This indicates a conformational change of the head domain away from the rest of the vimentin protein.

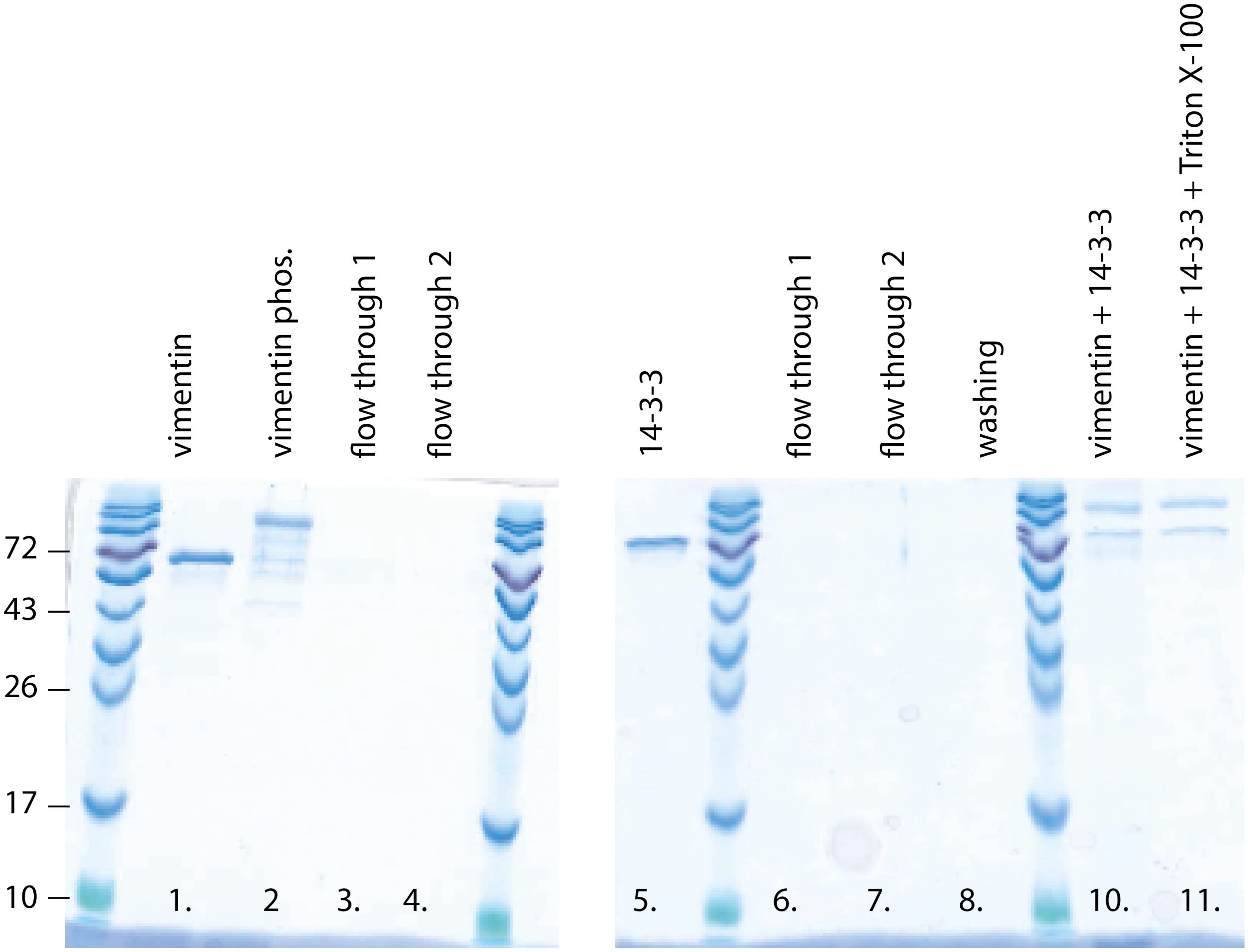


Figure 14: Pulldown assay shows interaction between vimentin and 14–3–3. This SDS gel shows the results of the streptavidin-biotin pull-down to investigate whether vimentin and 14–3–3 form a complex. The first lane shows vimentin and the second lane phosphorylated vimentin. Lane 3 and 4 show the flow through after incubation of the streptavidin agarose beads with vimentin. No vimentin is visible, indicating that the vimentin successfully bound to the beads. Lane 5 shows 14–3–3. Lane 6 and 7 show the flow through after the incubation of the beads bound to vimentin and 14–3–3. No band is observed, confirming the binding. Lane 8 shows a washing step in-between. Lane 10 and 11 show the protein bound to the beads with two bands at the weight of vimentin and 14–3–3. All unbound protein was washed off, vimentin bound to the beads *via* biotin-streptavidin binding and 14–3–3 formed a complex with the bound vimentin, therefore 2 bands are identified.

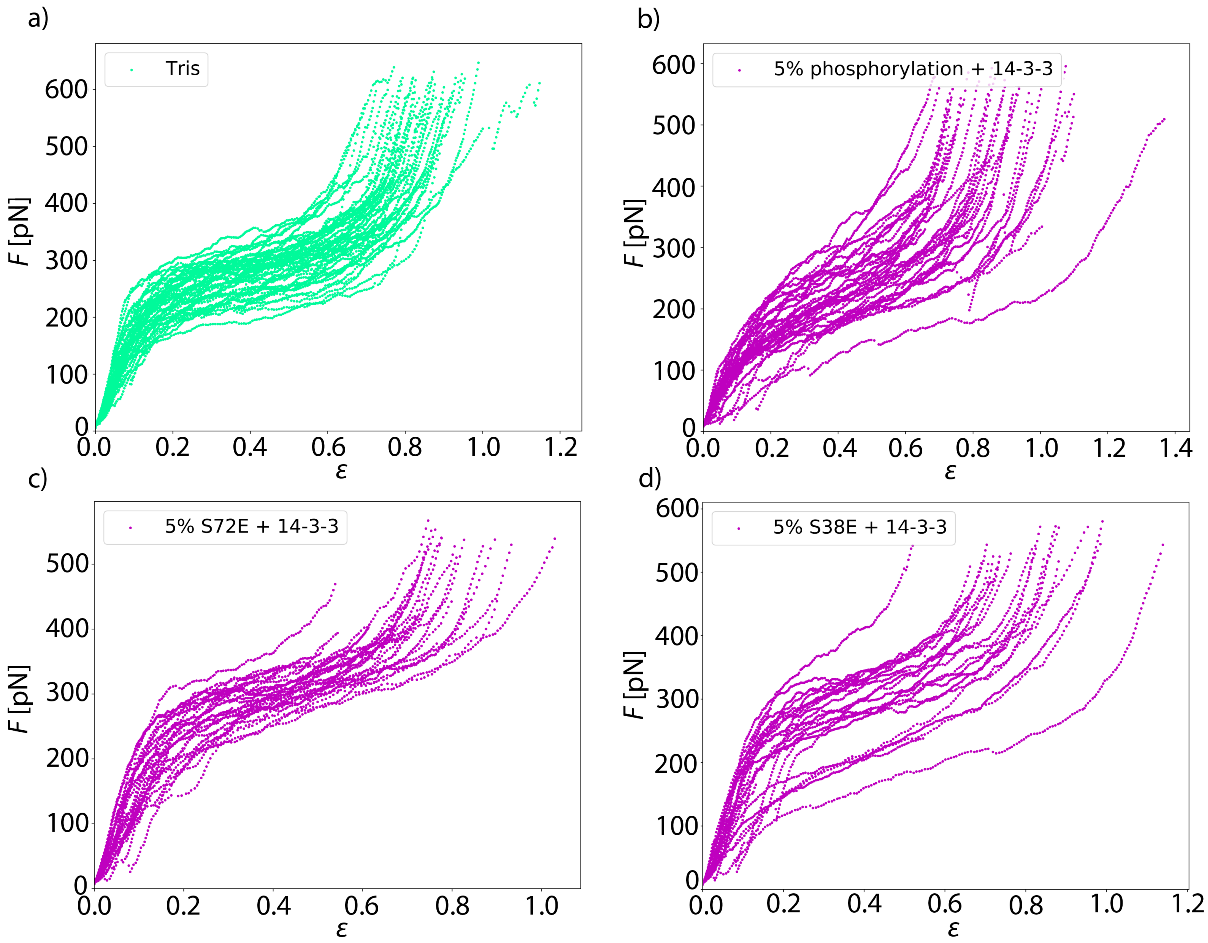


Figure 15: Single force-strain curves of all vimentin filaments incubated with 14–3–3. a) The control measurement without any modifications. b) Vimentin filaments with 5 % phosphorylated vimentin incubated with 14–3–3 shows softer filaments compared to the control. c) Vimentin filaments with 5 % of the S72E mutant incubated with 14–3–3 shows similar stiffness compared to the control. d) Vimentin filaments with 5 % of the mutant S38E also shows comparable stiffness to the control. The incubation of 14–3–3 with the mutations S72E and S38E does not change the force-strain behavior.

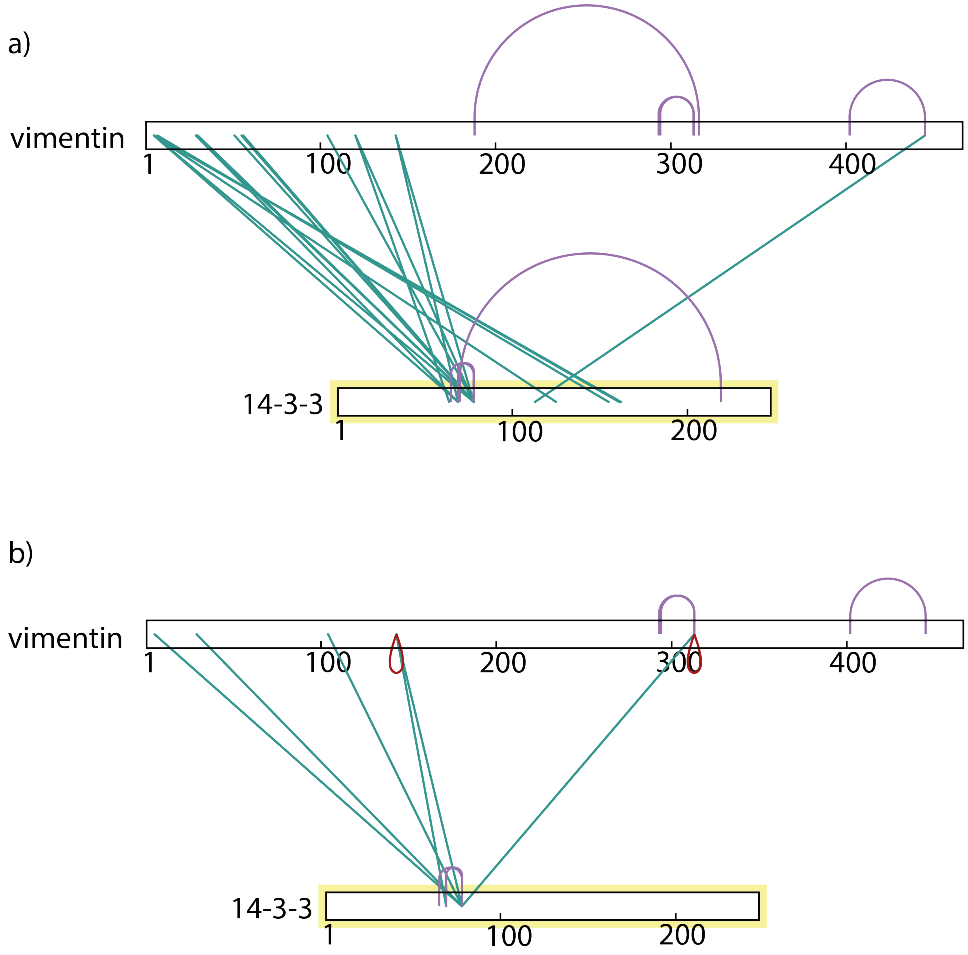


Figure 16: Mass spectrometry shows interaction sites of vimentin and 14–3–3. a) Vimentin monomer cross-linked with a 14–3–3 monomer. Green lines show cross-links between vimentin and 14–3–3, purple lines show cross-links within vimentin or 14–3–3 itself. The position that gets cross-linked most in the amino acid sequence of 14–3–3 is 78. It is linked to various positions in the amino acid sequence of vimentin, mostly found in the head region of vimentin. b) Vimentin dimer cross-linked with a 14–3–3 dimer. The color code of the lines is the same as above and red lines show cross-links within vimentin itself at the same position. The overall number of cross-links decreases, which is in accordance with limited sterical possibilities to form cross-links in a dimer compared to a monomer. The main cross-linking position in 14–3–3 is again 78 and two cross-linking positions (5, 29) for vimentin are located in the head domain of vimentin.

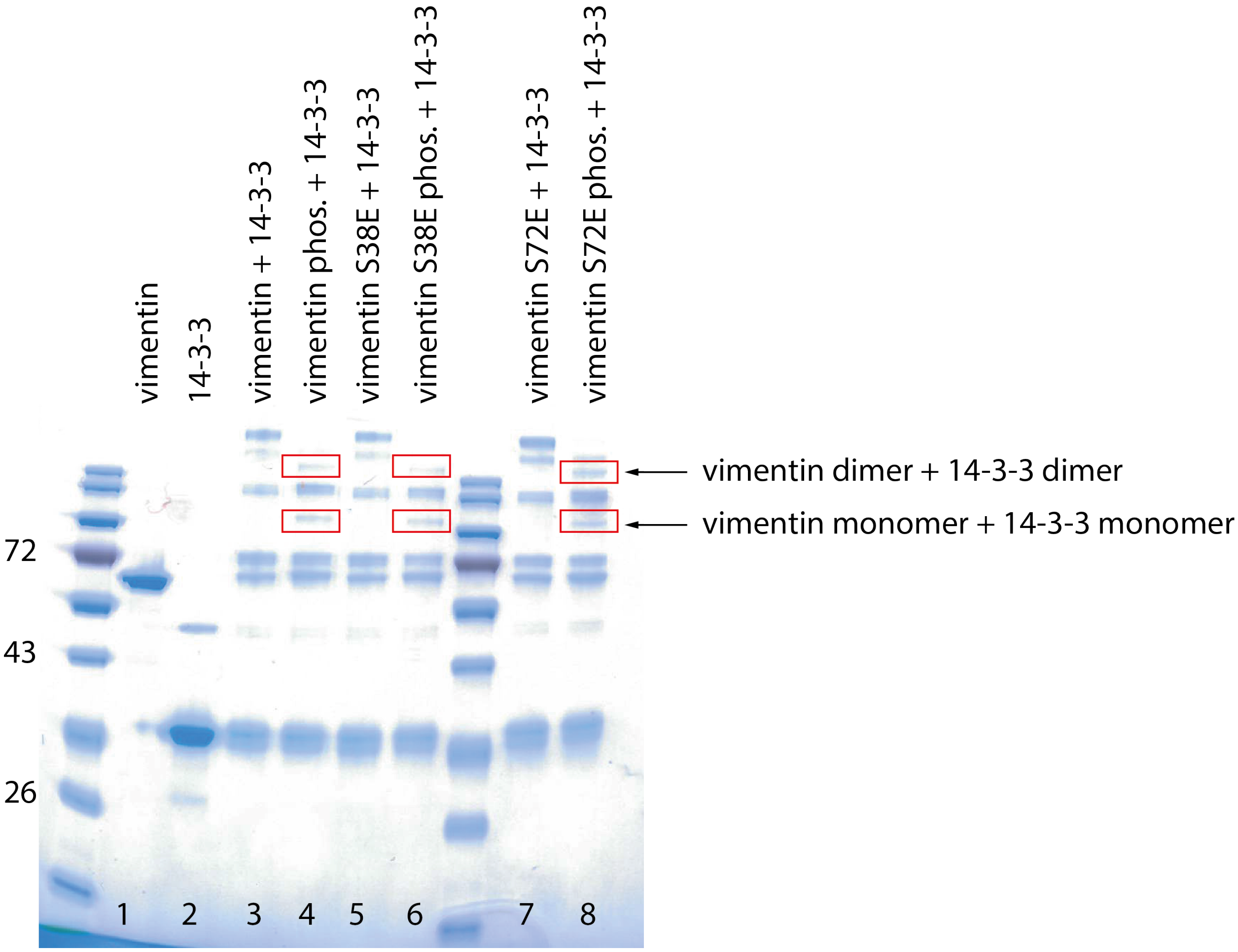


Figure 17: Investigation of the interaction between vimentin and 14–3–3 by cross-linking experiments. SDS gel of cross-linked probes. Cross-linking was performed with $BS^3$. Lane 1 shows the vimentin control with no cross-linkers and lane 2 the control for 14–3–3 without cross-linkers. In lane 3, vimentin cross-linked with 14–3–3 and in lane 4 phosphorylated vimentin cross-linked with 14–3–3 are shown. In lane 5 vimentin S38E cross-linked with 14–3–3 and in lane 6 phosphorylated vimentin S38E cross-linked with 14–3–3 are shown. In lane 7 vimentin S72E cross-linked with 14–3–3 and in lane 8 phosphorylated vimentin S72E cross-linked with 14–3–3 are shown. We observe that the phosphorylated form of the different vimentin types cross-linked with 14–3–3 results in additional bands (indicated by red boxes and arrows), which show that there is a protein complex. The phosphomimetic mutants in their unphosphorylated form (lane 5 and 7) do not interact with 14–3–3. Only for the phosphorylated form of vimentin complexes with 14–3–3. Furthermore the phosphomimetic mutants complex with 14–3–3, showing that the sites S38 and S72 are not the binding sites of 14–3–3.

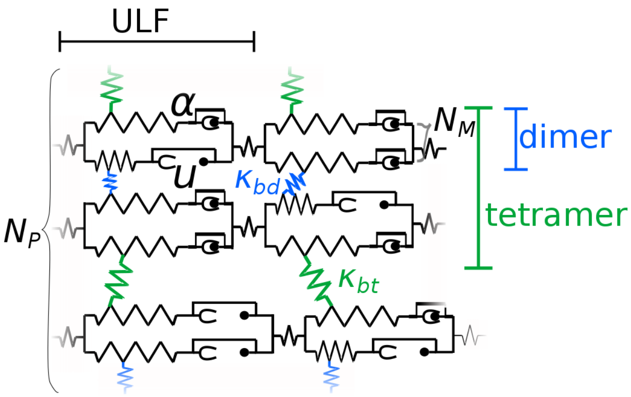


Figure 18: Sketch of the theoretical model for simulated force-strain curves. Each monomer is represented by a spring with a spring constant $\kappa_\alpha$ and by an element, which opens into an unfolded state $u$ once force is applied. Two monomers are connected in parallel to form a dimer, thus the number of monomers in the dimer is $N_M = 2$. The dimers are connected by additional springs (blue) with a spring constant $\kappa_{bd}$ to form tetramers. The connection between tetramers is established by another spring (green) with a spring constant $\kappa_{bt}$. Eight parallel tetramers ($N_P = 8$) form a ULF. The ULFs are connected in series *via* springs to a filament.

rmann, H.; Dadlez, M. Structural Dynamics of the Vimentin Coiled-coil Contact Regions Involved in Filament Assembly as Revealed by Hydrogen-Deuterium Exchange. *Journal of Biological Chemistry* **2016**, *291*, 24931–24950.

(5) Herrmann, H.; Häner, M.; Brettel, M.; O Ku, N.; Aebi, U. Characterization of distinct early assembly units of different intermediate filament proteins. *J. Mol. Biol.* **1999**, *286*, 1403–20.

(6) Mücke, N.; Wedig, T.; Bürer, A.; Marekov, L. N.; Steinert, P. M.; Langowski, J.; Aebi, U.; Herrmann, H. Molecular and biophysical characterization of assembly-starter units of human vimentin. *Journal of Molecular Biology* **2004**, *340*, 97–114.

(7) Kilisch, M.; Lytovchenko, O.; Arakel, E. C.; Bertinetti, D.; Schwappach, B. A dual phosphorylation switch controls 14-3-3-dependent cell surface expression of TASK-1. *J. Cell Sci* **2016**, *129*, 831–842.

(8) Block, J.; Witt, H.; Candelli, A.; Peterman, E. J. G.; Wuite, G. J. L.; Janshoff, A.;

Köster, S. Nonlinear Loading-Rate-Dependent Force Response of Individual Vimentin Intermediate Filaments to Applied Strain. *Physical Review Letters* **2017**, *118*, 1–5.

(17) Cox, J.; Mann, M. MaxQuant enables high peptide identification rates, individualized

ppb-range mass accuracies and proteome-wide protein quantification. *Nature Biotechnology* **2008**, *26*, 1367–1372.

(18) Tyanova, S.; Temu, T.; Cox, J. The MaxQuant computational platform for mass spectrometry-based shotgun proteomics. *Nature Protocols* **2016**, *11*, 2301.

(19) Consortium, U. UniProt: a worldwide hub of protein knowledge. *Nucleic Acids Research* **2019**, *47*, D506–D515.

(20) MacLean, B.; Tomazela, D. M.; Shulman, N.; Chambers, M.; Finney, G. L.; Frewen, B.; Kern, R.; Tabb, D. L.; Liebler, D. C.; MacCoss, M. J. Skyline: an open source document editor for creating and analyzing targeted proteomics experiments. *Bioinformatics* **2010**, *26*, 966–968.

(21) Konarev, P. V.; Volkov, V. V.; Sokolova, A. V.; Koch, M. H. J.; Svergun, D. I. PRIMUS: a Windows PC-based system for small-angle scattering data analysis. *J. Appl. Crystallogr.* **2003**, *36*, 1277–1282.

(22) Eigel, M.; Gruhlke, R.; Marschall, M.; Trunschke, P.; Zander, E. `alea` - A Python Framework for Spectral Methods and Low-Rank Approximations in Uncertainty Quantification. `https://bitbucket.org/aleadev/alea`.

(28) Kornreich, M.; Avinery, R.; Malka-Gibor, E.; Laser-Azogui, A.; Beck, R. Order and disorder in intermediate filament proteins. *FEBS Lett. 589*, 2464–2476.

(29) Herrmann, H.; Aebi, U. Intermediate Filaments: Strucutre and Assembly. *Cold Spring Harb. Perspect. Biol.* **2016**, *8*, a018242.
